# The chloroplast CLP chaperone-protease system controls the steady-state abundance of the singlet oxygen sensors EXCUTER 1 and 2

**DOI:** 10.64898/2026.08.05.743032

**Authors:** Claire M. Ravenburg, Pratyush Routray, Imen Bouchnak, Bingjian Yuan, Magdalena M. Julkowska, Klaas J. van Wijk

## Abstract

- The chloroplast CLP chaperone-protease is essential for chloroplast biogenesis. CLP substrate selection is aided by the N-recognin CLPS1 and CLPF adaptors. They interact with each other and the CLPC1 chaperone, but their specific functions are poorly understood.
- We employed *in vivo* CLPC1 substrate-trapping in Arabidopsis by expressing a *35S:CLPC1-TRAP-STREPII* transgene in wild-type (WT), *clpf, clps1,* and *clpfclps1* to test the consequences of the loss of these adaptors on CLPC1-trapped proteins. Immunoblotting and protein half-life experiments were carried out for identified CLP substrates.
- Expression of the *35S:CLPC1-TRAP-STREPII* in *clps1cpf* was embryo lethal. CLPF was trapped at a reduced level in *clps1*, supporting CLPS-CLPF interactions. Chloroplast ^1^O_2_ sensor EXECUTER1 (EX1) was trapped in WT and *clps1* but not significantly in *clpf*. Steady-state protein accumulation of EX1 and its homolog EX2 increased 30-fold in the CLPC1-TRAP lines and *clpr2-1*, but not in *clpf* or *clps1*. *In planta* experiments showed that the half-life of EX1 is ∼3-fold longer in *clpc1-1* than in WT, but EX1 half-life was unaffected in *clpf*.
- We conclude that the CLP system plays a key role in EX1,2 homeostasis by keeping their intra-chloroplast concentrations low through continuous degradation, upstream of their ^1^O_2_ signaling function.

## Introduction

Chloroplast protein homeostasis is established through regulated protein synthesis, import, folding, and degradation (Llamas & Pulido, 2022; Gao *et al*., 2023; Sun & Jarvis, 2023). These processes allow for chloroplast proteome remodeling in response to development, changing environmental conditions, and stress. Intra-chloroplast proteases and peptidases remove damaged, unfolded, or unwanted polypeptides, including the toxic cleaved chloroplast transit peptides, and recycle the products into free amino acids (Nishimura *et al*., 2017; van Wijk, 2024). Most chloroplast proteases have prokaryotic origins, arising from the endosymbiotic event that initiated chloroplast evolution, but these proteases have since diversified to support their unique roles in chloroplasts. While many chloroplast proteases have been characterized individually, little is known about their substrates and contributions to the overall chloroplast proteolysis network (van Wijk, 2024).

The CLP chaperone-protease system is the most abundant and complex protease in the chloroplast stroma, and the CLP system is essential for chloroplast biogenesis and plant development (Nishimura & van Wijk, 2015; Rodriguez-Concepcion *et al*., 2019; Bouchnak & van Wijk, 2021). The chloroplast CLP system in *Arabidopsis thaliana* is comprised of hexameric ATP-dependent AAA+ chaperones (CLPC1, CLPC2, and CLPD) that unfold substrates driven by ATP hydrolysis and feed them directly into a ∼350 kDa serine protease core complex for degradation into 7-11 amino acid peptides. The proteolytic core contains stacked heptameric P- and R-rings with subunits in a fixed stoichiometry. The R ring contains CLPP1, CLPR1-4 in a 3:1:1:1 ratio, while the P ring contains CLP3-6 with 1:2:3:1 copies, respectively (Olinares *et al*., 2011). Unlike the CLPP subunits, the CLPR subunits are all catalytically inactive; however, both loss-of-function CLPR and CLPP Arabidopsis mutants are seedling- or embryo-lethal, indicating that each core subunit is uniquely required for proper CLP function. CLPT1,2 proteins resembling N-domains of the CLPC1,2 chaperones associate with the CLP core, but their function is not clear (Sjogren & Clarke, 2011; Colombo *et al*., 2014; Kim *et al*., 2015).

Substrate unfolding by chloroplast CLP chaperones is required for substrate degradation by the chloroplast CLP protease system; this has also been extensively demonstrated for bacterial CLP chaperone-protease systems (Olivares *et al*., 2018; Mabanglo & Houry, 2022; Sauer *et al*., 2022). Mutations in the Walker B (ATPase) domains of the chaperones inhibit ATP hydrolysis, prevent substrate unfolding, and stabilize the interaction between CLPC and substrates, thereby enabling substrate identification (Rei Liao & van Wijk, 2019). This CLPC1 ‘trapping’ approach has successfully identified many confirmed and putative substrates of the Arabidopsis chloroplast CLP system (Montandon *et al*., 2019b; Rei Liao *et al*., 2022).

Substrate selection involves interaction with the N-domain of the CLP chaperones but can also be aided by adaptors, such as the N-recognin CLPS, as best demonstrated for bacterial CLP systems (Kuhlmann & Chien, 2017; Mahmoud & Chien, 2018). Arabidopsis contains a homolog of *E. coli* CLPS, CLPS1, which likely functions in an N-degron pathway in chloroplasts (Nishimura *et al*., 2013; Nishimura *et al*., 2015; Colombo *et al*., 2018; Montandon *et al*., 2019a; Aguilar Lucero *et al*., 2021; Kim *et al*., 2021). CLPS1 interacts with the CLPC1,2 chaperones, several confirmed CLP substrates such as GluTR, and the proposed co-adaptor CLPF (Nishimura *et al*., 2013; Nishimura *et al*., 2015). CLPF is distinct from other confirmed CLPS1 interactors because CLPF does not rely on the CLPS1 N-degron binding residues for interaction. Additionally, CLPS1 over-accumulates in *clpf* loss-of-function Arabidopsis plants, suggesting possible compensatory roles (Nishimura *et al*., 2015). CLPF also interacts with the CLPC1,2 chaperones not only at their N-domain, the canonical site of substrate binding, but also through their middle domains, which include a conserved UVR motif (Nishimura *et al*., 2015). Proteins that interact with the UVR motif of bacterial CLP chaperones often regulate CLP chaperone activity or oligomerization (Annis *et al*., 2024). Taken together, CLPF and CLPS1 are predicted to function as adaptors, anti-adaptors, and/or regulators of the chloroplast CLP system. However, the specific roles and mechanisms of CLPF and CLPS1 are unknown. Additionally, we do not yet know to what extent these adaptors act as a binary pair or if they also have independent functions.

In this study, we tested if CLP substrate selection is affected by the absence of the CLPF and CLPS1 adaptors by performing *in vivo* CLPC1 substrate-trapping in *clpf* and *clps1* loss-of-function Arabidopsis lines. Interestingly, the loss of CLPF, but not CLPS1, suppresses the yellow-heart leaf phenotype of CLPC1-TRAP observed in the WT background by reducing expression of the *CLPC1-TRAP* transgene. Previously, we showed that the chloroplast singlet oxygen response protein, EXECUTER1 (EX1), is trapped at high levels in WT/CLPC1-TRAP plants (Montandon *et al*., 2019b; Rei Liao *et al*., 2022). Here, we show that loss of CLPF but not loss of CLPS1 greatly reduced the amount of trapped EX1. Steady-state accumulation levels of EX1 and its homolog EX2 were low in WT plants but increased ∼30-fold in the *clpr2-1* knockdown and CLPC1-TRAP plants. *In vivo* half-life studies with a translational inhibitor show that the half-life of EX1 is significantly increased in CLP loss-of-function mutants. This strongly suggests that accumulation levels of EX1 and its homolog EX2 are kept low through continuous CLP degradation upstream of the ^1^O_2_ sensor function of EX1,2 and thylakoid FtsH triggered degradation and retrograde signaling.

## Material and Methods

### Plant Materials and Growth Conditions of Arabidopsis thaliana (Col-0)

In this study, we used the following previously established Arabidopsis (Col-0) lines: *clps1*, *clpc1-1, clpc1-1clps1* (Nishimura *et al*., 2013), *clpf, clps1clpf* (Nishimura *et al*., 2015), *clpr2-1* (Rudella *et al*., 2006), *ex1* (Wagner *et al*., 2004), and WT/*35S:CLPC1-TRAP-STREPII (Montandon et al., 2019b)*. We generated *clpfclpc1-1* by crossing and several additional CLPC1-TRAP lines as described below. Seeds were sown on agar (phytobend) plates with either Arabidopsis Germination Medium (Phytotech Labs) or ½ Murashige and Skoog (MS) medium without sucrose. Following a 2- or 3-day stratification at 4 °C, plates were transferred to light for 10-12 days, followed by transfer of the plantlets to soil. Plants were grown in a light/dark regime of 10 h light / 14 h dark (short day) at 22 °C, 60% relative humidity, and either 80 µmol photon s^-1^ m^-2^ (on plates) or 120 µmol photon s^-1^ m^-2^ (on soil). Plants were genotyped using genomic DNA extraction followed by PCR (30 cycles) and RNA extraction/cDNA synthesis followed by RT-PCR (22-25 cycles) using standard procedures and instructions by the manufacturers. The primers used are listed in Table S1.

### Generating the clpf/35S:CLPC1-TRAP-STREPII and clps1/35S:CLPC1-TRAP-STREPII Arabidopsis lines

Heterozygous WT/*35S:CLPC1-TRAP-STREPII* (Aa) plants were crossed with homozygous *clps1clpf (bbcc)* plants. We selected successfully crossed F1 plants based on the yellow-heart phenotype of heterozygous CLPC1-TRAP plants. Since *clpfclps1* plants do not have a visible phenotype, and the pollen donor for the cross was WT/*35S:CLPC1-TRAP-STREPII*, a virescent phenotype indicated a successful outcross rather than a self-cross of the *clpfclps1* plant. The *clps1* line (SAIL) and the *CLPC1-TRAP* transgenic plants are both BASTA-resistant, so we did not use BASTA selection media to confirm transgene presence in the F2 and F3 generations. Instead, we genotyped the F2 and F3 plants for *clps1* and *clpf* using gene-specific and TDNA primers, and for the presence of the *CLPC1-TRAP* transgene to obtain the *clpf/35S:CLPC1-TRAP-STREPII* and *clps1*/*35S:CLPC1-TRAP-STREPII* lines used in this study. The F3 and F4 generations of these lines were used for the remaining experiments. Zygosity of the transgene in the *clpf*/*35S:CLPC1-TRAP-STREPII* plants was determined by testing the segregation of the progeny on selection media (10 µg/mL BASTA). Seed abortions were quantified in developing (green) siliques in the F2 and F3 plants by removing the siliques from the inflorescence (3 siliques per individual), clearing the siliques in 70% EtOH for at least 48 hours, and manually counting the presence/absence of seeds under a light microscope. X^2^ analysis was performed in Excel. Additionally, independent WT/*35S:CLPC1-TRAP-STREPII* lines were created as described in (Montandon *et al*., 2019b) and crossed to *clpf* to verify the suppression of the CLPC1-TRAP phenotype by *clpf*.

### Photosynthetic Capacity Phenotyping

Arabidopsis seeds were sown on agar (phytobend) plates with Arabidopsis Germination Medium (Phytotech Labs). Following a 3-day vernalization at 4 °C, the plates were moved to a chamber with a 16-hour light/8-hour dark photoperiod, light intensity of 150 µmol photons s^-1^ m^-2^, 20/18 °C day/night temperature, and 60% relative humidity. After 10 days, the plantlets were transplanted to soil. Following a 2-day recovery period, the plants were moved to and monitored in the PhenoSight Facility at the Boyce Thompson Institute under controlled environmental conditions (16-hour light/8-hour dark photoperiod, light intensity of 150 µmol photons s^-1^ m^-2^, 20/18 °C day/night temperature, and 60% relative humidity). Plants were maintained at a target pot weight of 130 g, corresponding to approximately 60% of soil water-holding capacity. Imaging was performed every 12 h—once near the end of the dark period and once near the end of the light period. Before each imaging session, plants were dark-adapted for 15 min in a dedicated adaptation tunnel. Chlorophyll fluorescence parameters were recorded using the CropReporter system (PhenoVation B.V.) following the pulse amplitude modulation (PAM) method. Minimum (F₀) and maximum (Fₘ) fluorescence were measured in the dark-adapted state, followed by a 30 sec light adaptation period to determine the light-adapted minimum (F₀′) and maximum (Fₘ′) fluorescence. After chlorophyll fluorescence measurements, plants were transferred to the RGB/ChlF imaging station, where top- and side-view images were collected.

### Affinity Purification of CLPC1-TRAP-STREPII

Batches (10 g each) of mature (stage 5.10) Arabidopsis rosettes from *35S:CLPC-TRAP-STREPII* lines in WT, *clps1* and, *clpf* backgrounds, were ground by mortar and pestle in liquid nitrogen to a fine powder, transferred to a tube, and vortexed in 1 mL extraction buffer (EB: 50 mM HEPES-KOH pH 7.8, 15% glycerol, 10 mM MgCl_2_, 75 mM NaCl, 250 μg/mL avidin, and 250 μg/mL Pefabloc serine protease inhibitor) per 1 g of fresh weight. The suspension was filtered through four layers of Miracloth (25 μm, Millipore) followed by centrifugation for 1.5 hours at 24,300 rpm in a SW27 rotor at 4 °C. The supernatants (input) were collected and either directly used for affinity purification on StrepTactinXT 4Flow affinity resin (IBA Life Sciences) or stored at −80 °C for later analysis. StrepTactin columns (column volume (CV): 1 ml) were prepared and washed with 2 CVs of Wash Buffer without glycerol (WB: 50 mM HEPES-KOH pH 7.8, 15% glycerol, 10 mM MgCl_2_, 75 mM NaCl), followed by equilibration with 2 CV of WB. The inputs were then applied to the column, followed by column washing with 10 CV of WB. STREPII-tagged proteins were eluted in 4 CVs of WB and 50 mM biotin. The eluates were pooled and concentrated using Ultra-4 Centrifugal Filter Units with a 3-kDa cutoff by centrifugation for about 3 hours at 7,500 xg at 4 °C in a JA25.15 rotor, resulting in a final volume of 50-70 μL. The concentrates were aliquoted and stored at −80 °C for MS/MS and immunoblot analysis. Two completely independent experiments were carried out (different sets of plants grown at different times of the year). Each experiment consisted of 3 biological replicates per genotype (purifications from which were completed on 3 consecutive days). Between purifications of biological replicates, the columns were regenerated with 15 CVs of 3 M MgCl_2_, then washed with 8 CVs of WB without glycerol.

### MSMS sample preparation, data acquisition, processing and analysis

Affinity eluates from the CLPC1-TRAP-STREPII purifications were separated by SDS-PAGE on Biorad Criterion Tris-HCL precast gels (10.5-14% acrylamide gradient) (Fig. S3). The gels were fixed with 5% AcOH and 50% EtOH, stained with 0.04% Coomassie Blue R250 (dissolved in fix solution), destained with 5% AcOH and 30% EtOH, and stored in 1% AcOH. After washing the gel twice for 5 minutes with diH_2_O, each gel lane was cut into consecutive gel slices (7 per lane), followed by reduction, alkylation, and in-gel digestion with trypsin as described in (Friso *et al*., 2011).

Extracted peptides were resuspended in 2% formic acid and analyzed using a QExactive mass spectrometer (ThermoElectron) equipped with a nanospray flex ion source and interfaced with a nanoLC system and autosampler (Dionex Ultimate 3000 Binary RSLCnano system) as described in (Rei Liao *et al*., 2022) with the exception that AGC target values were set at 1 x 10^6^ for the MS survey scans and maximum scan time 30 ms and 5.10^5^ for MSMS scans and maximum scan time 50 ms. Other parameters for MS were 70k resolution, scan range 400-2000 m/z, and for MSMS were 17.5k resolution, top10, loop count 10, isolation window 2.0 m/z, dynamic exclusion 15 sec.

Peak lists in MGF format were generated from RAW files using Distiller software (version 2.7.1.0) in default mode (Matrix Science). MGF files were searched with MASCOT v2.4.0 against TAIR10 including a set of typical contaminants and the decoy (71148 sequences; 29099536 residues). Two parallel searches (Mascot p-value <0.01 for individual ion scores; precursor ion window 700 to 3500 Da) were carried out: (i) Full tryptic (error tolerance 10 ppm for MS and 0.5 Da for MSMS) and (ii) Semi-tryptic (error tolerance 6 ppm and 0.5 Da for MSMS) in both cases with fixed Cys-carbamido-methylation and variable M-oxidation, N-terminal acetylation, N-terminal formylation and 2 missed cleavages (in Mascot PR or PK does not count as missed cleavage). In-house post-Mascot filtering removed redundancies between the full and semi-tryptic searches and removed spectral matches with ion scores below 33. The semi-tryptic search served to increase protein coverage and was combined with the full tryptic search results. Proteins identified by MSMS spectra that were all shared with other proteins identified by unique peptides were discarded. Final false discovery rate for proteins identified with one spectral match was below 1% and zero for those with two or more spectral matches.

Proteins were quantified by the spectral counting method (SPC) using matched peptide spectra. For quantification by spectral counting, each protein accession was scored for total spectral counts (SPC), unique SPC (uniquely matching to an accession), and adjusted SPC (Friso *et al*., 2011). The latter assigns shared peptides to protein accessions in proportion to their relative abundance using unique spectral counts for each accession as a basis. After selecting only the best-scoring isoform model per protein, proteins that shared a significant number of matched spectra were grouped, and proteins and groups with less than 3 adjSPC were discarded for further analysis.

For significance analysis and volcano plots to compare the *clps1* and *clpf* backgrounds with WT backgrounds, a minimal total adjSPC count of 12 was required to ensure a more robust dataset and lower FDR. After imputation of the value 0.99 for the adjSPC of protein accessions with missing values within a replicate (a complete gel lane), the adjSPCs were normalized to the total adjSPC within a replicate, resulting in NadjSPC for each protein. Average NadjSPC per genotype across the six replicates was calculated, and the p-values were determined by the student t-test to determine proteins that showed significant pairwise abundance differences.

### Total and Soluble Leaf Protein Extractions for Immunoblot Analysis

Arabidopsis seedlings (stage 1.04-1.06) were harvested from agar plates, roots were removed, and the remaining leaf tissue was immediately flash frozen in liquid nitrogen. Seedlings were ground using a TissueLyser (Qiagen) and zinc-plated 4.5 mm BBs for 40 seconds at 30 rpm. 2 µL of either total protein extraction buffer (50 mM Tris-HCl (pH 8), 15% glycerol, 2% SDS, and 250 µg/mL Pefabloc) or soluble protein extraction buffer (50 mM HEPES-KOH pH 7.8, 15% glycerol, 10 mM MgCl_2_, 75 mM NaCl, and 250 µg/mL Pefabloc) per 1 mg of fresh tissue was added to the tissue powder before vortexing the sample for 1 minute (resting on ice every 20 seconds). For soluble protein extractions, the samples were immediately centrifuged at 20,627 xg for 15 minutes at 4 °C to separate total soluble protein from membranes, aggregates, and starch granules. For total protein extractions, samples were incubated on ice for 5 minutes to ensure full solubilization of membranes, heated at 37 °C for 15 minutes, then centrifuged at 20,627 xg for 15 minutes at 4 °C. The soluble protein or total protein supernatants were then aliquoted and stored at -80 °C for analysis. Protein concentration of the samples was determined using the BioRad DC assay.

Aliquots of either total leaf protein, soluble leaf protein, or affinity-purified proteins were separated on Tris-HCl mini gels (6% acrylamide stacking, 12% acrylamide resolving), transferred to 0.2 µm nitrocellulose membranes (BIO-RAD), stained with Ponceau S, and analyzed by immunoblotting using chemiluminescence for detection. The antisera used were: anti-STREPII (1:2500 dilution; GenScript #10498-120), anti-EX1 and anti-EX2 (1:1000 dilution; PhytoAB #3099A and #1694, respectively), and sera produced by the van Wijk lab against purified recombinant proteins: anti-CLPF (1:2500 dilution) and anti-CLPS1 (1:1000 dilution) (Nishimura *et al*., 2015). Protein bands were quantified using ImageJ, and the resulting data were analyzed and graphed in Excel.

### RNA extraction and RT-PCR and qRT-PCR assays

Total RNA was isolated with the RNeasy plant mini kit (Qiagen) and treated with on-column DNase (Qiagen). cDNA was synthesized from equal amounts of total RNA with Superscript III Reverse Transcriptase following the manufacturer’s protocol using the provided Oligo(dT)_20_ primers (Invitrogen). Two or three biological replicates (4 seedlings per replicate) were analyzed for each experiment. Transcript levels of *EX1, CLPF, CLPS1, CLPC1-TRAP-STREPII* and *ACTIN2* (*ACT2*) were determined using RT-PCR. The reaction mixture (20 µL) contained the template (0.2-0.5 µg), 0.2 µM of each primer, and Promega GoTaq G2 Green Master Mix. The amplification conditions were as follows: 95 °C for 5 min, followed by 22-25 cycles of 95 °C for 30 s, 51-55 °C for 30 s, and 72 °C for 1 min/1kb amplicon, and a final extension step at 72 °C for 5 min. The results were normalized using ACT2 as the endogenous control. To analyze CLPC1-TRAP-STREPII and total CLPC1 transcript levels in the different CLPC1-TRAP lines, qRT-PCR was performed with PowerTrack SYBR Green Master Mix with three technical replicates (10 ng cDNA) per reaction. All qRT-PCR experiments were conducted with a QuantStudio 7 Pro qPCR machine, and the results were normalized to the geometric mean of *ACTIN2* (*ACT2* – AT3G18780) and *UBIQUITIN10* (*UBQ10* – AT4G05320) levels. All primer sequences are listed in Table S1.

### CHX chase assays for protein half-life analysis

Plant material for cycloheximide (CHX) chase assays to determine the half-life of EX1 was generated by germinating seeds and growing the resulting seedlings for 15-20 days in 10-hour light/14-hour dark at 22 °C, 60% relative humidity, and 80 µmol photon s^-1^ m^-2^ on ½ Murashige and Skoog (MS) agar plates without sucrose. The seedlings (including roots) at developmental stage 1.04-1.06 were then transferred to 12-well culture plates with 4 mL of liquid ½ MS media and incubated for two hours under the same light conditions as during growth with gentle shaking. After two hours, the liquid medium was supplemented with 300 µM CHX in DMSO or only DMSO for mock controls. Transpiration rates were increased by removing the lids from the culture plates and increasing air circulation with a fan. The seedlings were collected at different time points (∼20 mg corresponding to 5-6 seedlings per replicate; 3 biological replicates per time point), roots were removed, and the shoots were briefly dried to remove excess media before being frozen in liquid nitrogen and stored at -80 °C for total protein extraction and immunoblot analysis. 4–20% Criterion™ TGX™ Precast Midi Protein Gels with 26 wells (BIO-RAD) were used.

## Results

### Generation of 35S:CLPC1-TRAP-STREPII lines in clps1, clpf, and clps1clpf null lines

We generated *clpf/35S:CLPC1-TRAP-STREPII* and *clps1*/*35S:CLPC1-TRAP-STREPII* Arabidopsis plants by crossing heterozygous WT/*35S:CLPC1-TRAP-STREPII* plants with homozygous *clps1clpsf* plants (Nishimura *et al*., 2015) (Fig. 1A). Crossing was used to avoid possible positional effects caused by introducing the *35S:CLPC1-TRAP-STREPII* transgenes into the different genetic backgrounds through floral dipping. Successful crosses (F1) were identified by the virescent phenotype caused by accumulation of CLPC1-TRAP-STREP protein (Montandon *et al*., 2019b) (Fig. 1B). In the segregating F2 and F3 generations, we then identified transgenic *CLPC1-TRAP-STREP* lines in homozygous *clps1* and homozygous *clpf* backgrounds through genotyping and RT-PCR (Fig. S1). The *clps1*/*35S:CLPC1-TRAP-STREPII* (Aa) seedlings have a yellow-heart phenotype, while the *clps1*/*35S:CLPC1-TRAP-STREPII* (AA) seedlings have albino/light green phenotypes (Fig. 1B); these albino seedlings never produce seeds, and they have poor viability when transferred to soil (data not shown). We note that the growth plates in Figure 1B contained 1% sucrose; in the absence of sucrose, these albino/light green seedlings are even smaller and white (data not shown). These phenotypes in the *clps1* background are similar to those when *35S:CLPC1-TRAP-STREPII* (AA or Aa) is expressed in the WT background (Fig. 1B). Surprisingly, *clpf*/*35S:CLPC1-TRAP-STREPII* (Aa) seedlings are visually nearly indistinguishable from WT plants, whereas *clpf*/*35S:CLPC1-TRAP-STREPII* (AA) seedlings have pale cotyledons, but leaf phenotypes similar to WT leaves (Fig. 1B). Additional phenotyping of plate-grown seedlings after transfer to soil further demonstrated the suppression of these *35S:CLPC1-TRAP* phenotypes in the *clpf* background (Fig. 1C). Figure 1D shows fully established rosette plants on soil of WT/3*5S:CLPC1-TRAP-STREPII* (Aa), *clps1*/3*5S:CLPC1-TRAP-STREPII* (Aa), *clpf/35S:CLPC1-TRAP-STREPII* (AA) demonstrating identical visible phenotypes (a developmental leaf gradient of yellow-pale green to full green) of these heterozygous transgenic lines in WT and *clps1*, but near WT-like green rosettes in the homozygous transgenic line (AA) in *clpf*. These three *35S:CLPC1-TRAP-STREPII* lines accumulate similar levels of CLPC1-TRAP protein as determined by immunoblotting against the STREPII tag, and immunoblotting confirmed the complete loss of CLPS1 and CLPF in *clps1* and *clpf,* respectively (Fig. 2A,B). Accumulation of CLPS1 increased 2-3-fold in *clpf*/*CLPC1-TRAPII*, similar to what was previously observed for *clpf* (Nishimura et al 2015) (Fig. 2A,B). This shows that CLPS1 accumulation level increases upon loss of CLPF, irrespective of the presence of CLPC1-TRAP protein, whereas the level of CLPF does not change in *clps1* backgrounds (Fig. 2A,B). Collectively, the data presented above show that loss of CLPS1 has no visible impact on the *35S:CLPC1-TRAP-STREPII* phenotypes, whereas loss of CLPF strongly suppressed the *35S:CLPC1-TRAPII* leaf phenotypes.

**Fig. 1.**
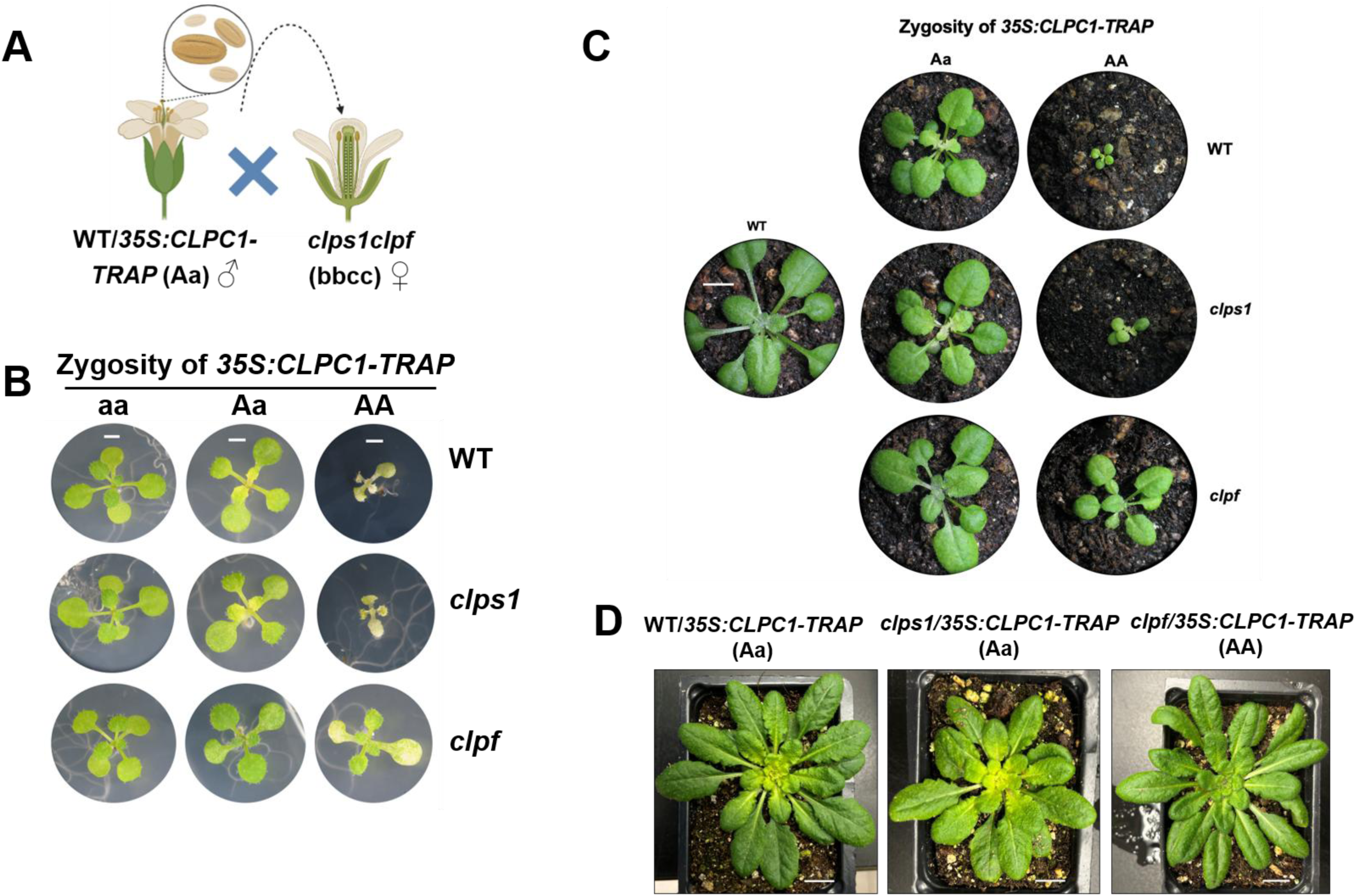
Generation and characterization of the *clps1*/*35S:CLPC1-TRAP-STREPII* and *clpf*/*35S:CLPC1-TRAP-STREPII* Arabidopsis lines**. A.** WT/*35S:CLPC1-TRAP-STREPII* (Aa) plants (the pollen donor) were crossed with *clps1clpf* (bbcc) plants, allowing for easy visual selection of successful crosses because expression of heterozygous *35S:CLPC1-TRAP-STREPII* results in a yellow-heart phenotype. **B**. Representative images of the phenotypic differences in 10-day-old WT, *clps1* (bb), and *clpf* (cc) Arabidopsis seedlings on ½ MS medium with 1% sucrose segregating for the *35S:CLPC1-TRAP-STREPII* transgene (aa, Aa, or AA). Scale bar is 1 mm. **C.** Phenotypes of 3-week-old WT/*35S:CLPC1-TRAP-STREPII* (Aa or AA), *clps1/35S:CLPC1-TRAP-STREPII* (Aa or AA), *clpf/35S:CLPC1-TRAP-STREPII* (Aa or AA), and WT on soil. Scale bar is 5 mm. **D.** Representative images of 6-week-old WT/*35S:CLPC1-TRAP-STREPII* (Aa), *clps1/35S:CLPC1-TRAP-STREPII* (Aa), *clpf/35S:CLPC1-TRAP-STREPII* (AA) on soil. Scale bar is 1 cm.

**Fig. 2.**
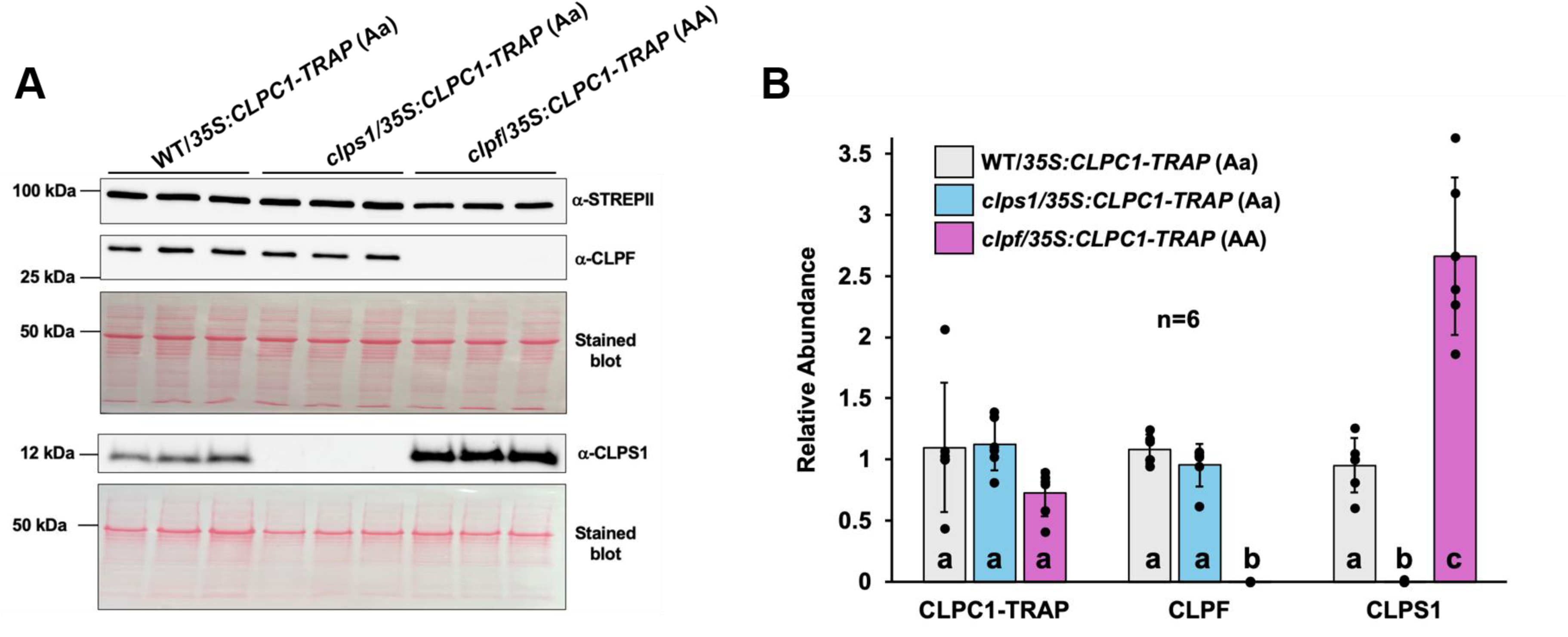
Steady state accumulation of CLPC1-TRAP, CLPS1, and CLPF in WT/35S:CLPC1-TRAP-STREPII (Aa), clps1/35S:CLPC1-TRAP-STREPII (Aa), and clpf/35S:CLPC1-TRAP-STREPII *(AA).* **A**. Representative immunoblots of total soluble leaf proteome for CLPC1-TRAP, CLPS1, and CLPF using specific antisera as indicated. Plants were at developmental stage 5.10 and were grown in soil for about 4 weeks after transplanting from plates. Corresponding Ponceau stained blots show the equal loading across samples. **B**. Bar diagram of the quantified CLPC1-TRAP, CLPF, and CLPS1 immunoblot signals relative to the WT/*35S:CLPC1-TRAP*-STREPII (Aa) sample in the WT/*35S:CLPC1-TRAP-STREPII* (grey bars), *clps1*/*35S:CLPC1-TRAP-STREPII* (blue bars), *clpf*/*35S:CLPC1-TRAP-STREPII* (purple bars) with n=6 (MS replicates 1-6). CLPS1 overaccumulates by ∼3-fold in the *clpf*/*35S:CLPC1-TRAP-STREPII* line compared to the WT/*35S:CLPC1-TRAP-STREPII* line. Significance was calculated using Dunnett’s multiple two-way ANOVA comparison (b,c = p-value < 0.001).

To verify that this suppression by *clpf* was not due to a positional effect of the transgene, we generated three independent *35S:CLPC1-TRAP-STREPII* transgenic lines. These new lines showed the same phenotypes as the original *35S:CLPC1-TRAP-STREPII* transgenic line (Fig. 3A; Fig. S2). Crossing these new lines to *clpf* null plants resulted in *clpf*/*35S:CLPC1-TRAP-STREPII* lines with similar suppression to the original line generated by the cross as illustrated for line #6 (Fig. 3B). Immunoblotting showed that the level of CLPC1-TRAP in both the original and independent *clpf*/*35S:CLPC1-TRAP-STREPII* (AA) lines is very similar as in the heterozygous *35S:CLPC1-TRAP-STREPII* lines in the WT and *clps1* backgrounds (Fig. 3C). qRT-PCR of the heterozygous *35S:CLPC1-TRAP-STREPII* mRNA in WT, *clps1,* and *clpf* showed that *CLPC1-TRAP* mRNA levels are reduced by ∼50% in the *clpf* background, compared to WT and *clps1* backgrounds. Analysis of the homozygous 35S:CLPC1-TRAP-STREPII lines in *clpf* showed a doubling of *CLPC1-TRAP* mRNA compared to these heterozygous lines in *clpf* (Fig. 3D). Previously we showed that homozygosity for the *35S:CLPC1-TRAP* gene doubles the amount of accumulated CLPC1-TRAP protein compared to the heterozygous line (Montandon *et al*., 2019b). Therefore, the suppression of the *35S:CLPC1-TRAPII* phenotype in the *clpf* background can, at least in part, be explained by reduced CLPC1-TRAP protein accumulation. The total mRNA levels for *CLPC1-TRAP* and endogenous CLPC1 showed only statistically significant increases in the homozygous *35S:*CLPC1-TRAP line in *clpf* as compared to wt and the other tested lines (Fig. 3E). To further capture and quantify the phenotypic differences between WT/*35S:CLPC1-TRAP-STREPII* (Aa) and *clpf*/*35S:CLPC1-TRAP-STREPII* (AA), we employed an automated imaging phenotyping facility to track plant growth and photosynthetic capacity of these lines, as well as *clpf* and WT plants (Fig. 4). Continuous imaging of the projected rosette area (top view) (Fig. 4A) showed no differences between WT and *clpf*, a small and consistent difference between WT and *clpf/35S:CLPC1-TRAP-STREPII (AA)* and large, persistent difference between WT and *clps1/35S-CLPC1-TRAP-STREPII* (Aa). Significant (p<0.05) differences in maximum quantum yields of photosystem II between WT and *clpf/35S:CLPC1-TRAP-STREPII* (AA) and between WT and WT*/35S:CLPC1-TRAP-STREPII* (Aa) were observed until after 14-21 days (350-400h) of imaging (Fig. 4B). Together, these observations show the *clpf* background strongly but not entirely suppresses the *35S:CLPC1-TRAP-STREPII* growth and photosynthetic phenotype, and that these phenotypes are strongest earlier in leaf and chloroplast development.

**Fig. 3.**
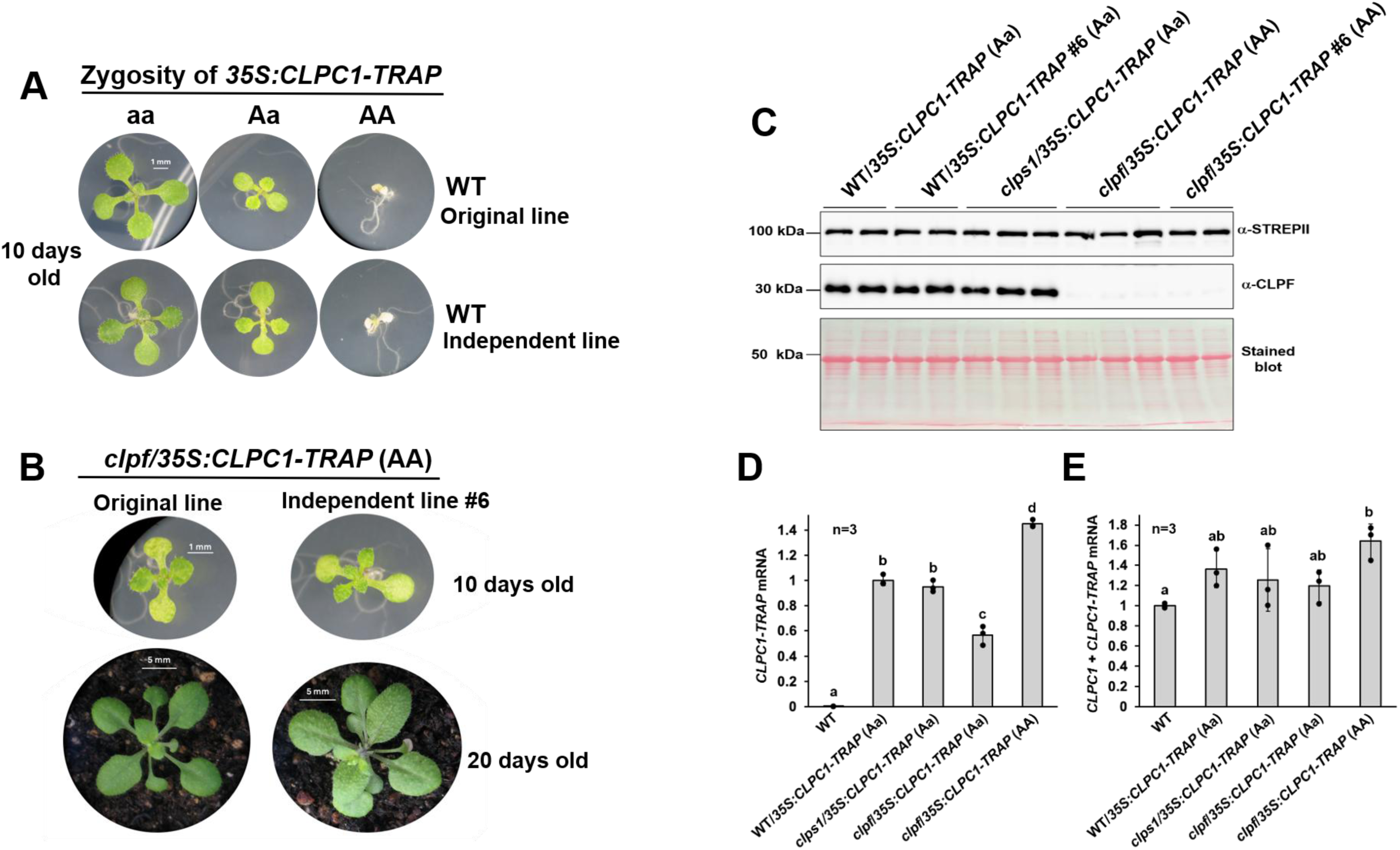
Characterization of an independent WT/*35S:CLPC1-TRAP-STREPII* and *clpf*/*35S:CLPC1-TRAP-STREPII* line and expression of *CLPC1-TRAP* and *CLPF*. **A**. Phenotypes of segregating progeny from a WT/*35S:CLPC1-TRAP-STREPII* (Aa) plant and an independent transgenic event WT/*35S:CLPC1-TRAP-STREPII* (Aa) #6. Both lines have a yellow-heart phenotype or albino/pale green phenotype when heterozygous or homozygous for the transgene, respectively. Seedlings were grown for 10 days on ½ MS without sucrose. **B.** Heterozygous WT/*35S:CLPC1-TRAP-STREPII* #6 (Aa) was crossed with *clpf* to test whether the loss of CLPF suppresses the phenotype in an independent *35S:CLPC1-TRAP* line. Indeed, *clpf*/*35S:CLPC1-TRAP-STREPII* #6 (AA) plants lack the conical albino phenotype at both developmental stages 1.04 and 1.10. **C.** Immunoblots of total soluble leaf proteome showing accumulation of CLPC1-TRAP and CLPF using specific antisera as indicated in five different genotypes as indicated. Corresponding Ponceau-stained blots show equal loading across samples. Plants were about 3 weeks old (developmental stage 1.08-1.10) **D,E.** qRT-PCR analysis of *CLPC1-TRAP-STREPII* and total *CLPC1* (endogenous + transgenic) transcript levels in WT, WT/*35S-CLPC1-TRAP-STREPII* (Aa), *clps1*/*35S:CLPC1-TRAP-STREPII* (Aa), and *clpf*/*35S:CLPC1-TRAP-STREPII* (AA and Aa) plants (stage 1.08-1.06). The data are normalized to both the *ACTIN2* and *UBIQUITIN1* levels of each sample. *CLPC1* transcript levels are shown relative to those in the WT samples, while *CLPC1-TRAP-STREPII* transcript levels are shown relative to those in the WT/*35S:CLPC1-TRAP-STREPII* (Aa) samples. Significance was calculated using Dunnett’s multiple two-way ANOVA comparison. In panel D, b,c,d = p-value < 0.0001, and in panel E, b = p-value < 0.05. n=3.

**Fig. 4.**
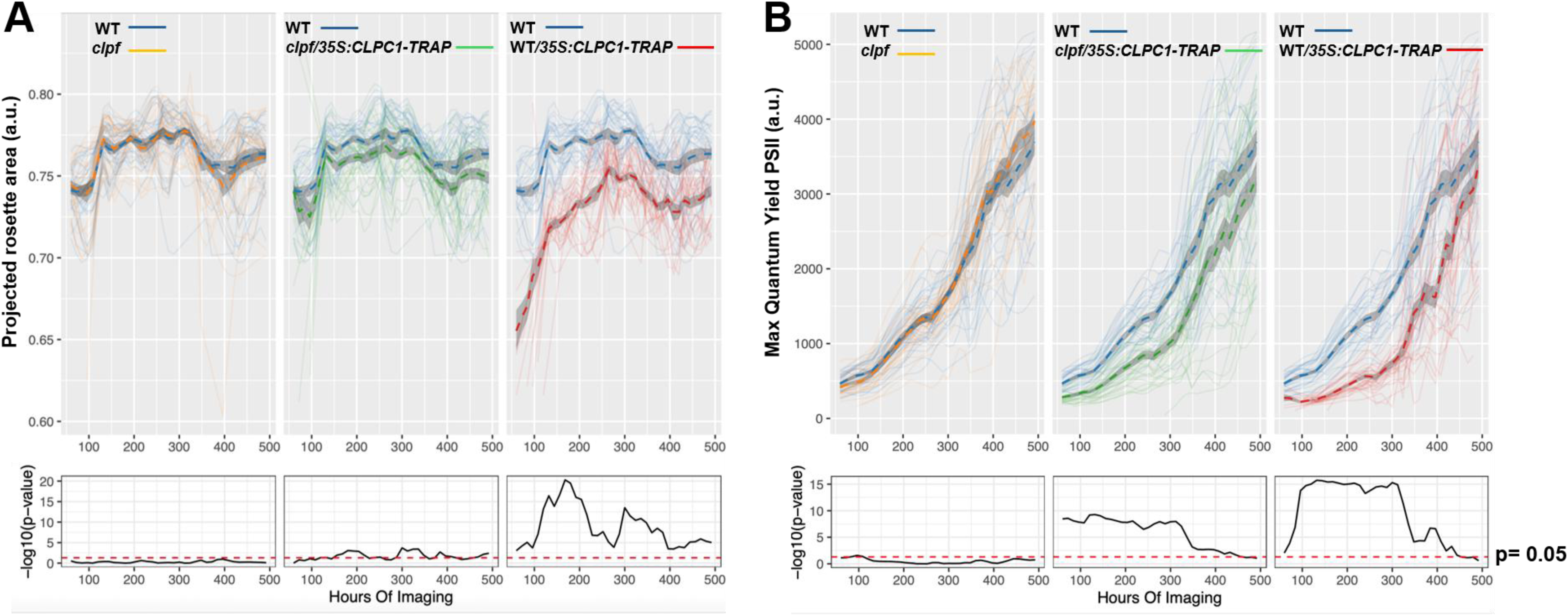
Automated phenotyping of soil-grown WT, *clpf*, WT/*35S:CLPC1-TRAP-STREPII* (Aa), and *clpf*/*35S:CLPC1-TRAP-STREPII* (AA). Plants were grown on ½ MS + 1% sucrose plates for 10 days. Plants were then imaged from day 12 (developmental stage 1.04-1.06) to day 34 (developmental stage 6.00). **A.** Projected rosette area. **B.** Dark-adapted Fv/Fm (maximum quantum yield of photosystem II). The calculated p-values (as -Log10 values) for the difference between WT and *clpf*, WT/*35S:CLPC1-TRAP-STREPII* (Aa), or *clpf/35S:CLPC1-TRAP-STREPII* (AA) are shown in the lower panels.

### Expression of CLPC1-TRAP-STREPII in the clps1clpf double mutant is embryo lethal

Surprisingly, we were unable to recover any *CLPC1-TRAP-STREPII* transgenic lines in the homozygous *clpfclps1* background; this was surprising because the *clpfclps1* double mutant has no visible growth or developmental phenotypes. Microscopy analysis of seeds in the developing siliques of *clps1*(Bb)*clpf*(cc)/*35S:CLPC1-TRAP-STREPII* (AA) plants showed a 1:3 ratio of shriveled, white inviable seeds and green seeds (Table 1; Fig. S3). Similarly, *clps1*(bb)*clpf*(Cc)/*35S:CLPC1-TRAP-STREPII* (Aa) plants (note that this line is heterozygous for the transgene) showed a 4.3:1 ratio of shriveled, white inviable seeds and green seeds (Table 1). This demonstrated that expression of *CLPC1-TRAP-STREPII* (AA or Aa) is embryo lethal in the *clpfclps1* background. We therefore continued the experiments with the *clpf*/CLPC1-TRAP-STREPII (AA) and the *clps1*/*35S:CLPC1-TRAP-STREPII* (Aa) lines.

**Table 1.** Quantification of the number of seed abortions observed in several siliques produced by F3-generation progeny of the *clps1clpf* (bbcc) x WT/*35S:CLPC1-TRAP-STREPII* (Aa) cross. A Chi-square test of ten F3 parents with a CLPS1*clpf*(Bbcc)/*35S:CLPC1-TRAP-STREPII* (AA) genotype shows that the viable-seed and embryo-aborted phenotypes segregate in a 3:1 ratio. χ2 (N = 396) = 0.9496, P > 0.05 supporting that the AAbbcc genotype is lethal. Likewise, a Chi-square test of five F3 parents with a *clps1*CLPF(bbCc)/*35S:CLPC1-TRAP-STREPII* (Aa) genotype shows that the viable-seed and embryo-aborted phenotypes segregate in a 13:3 ratio. χ2 (N = 226) = 0.2093, P > 0.05 supporting that both the Aabbcc and AAbbcc genotypes are lethal. Values in the table are the average number of viable seeds and abortions observed in three siliques per individual parent.

| Heterozygous<br>for *clps1*<br>(AABbcc) | Seeds in developing siliques |  |  | Heterozygous<br>for *clpf*<br>(AabbCc) | Seeds in developing siliques |  |  |
| --- | --- | --- | --- | --- | --- | --- | --- |
| parent # | alive | dead | total | parent # | alive | dead | total |
| 1 | 30 | 8 | 38 | 1 | 32 | 11 | 43 |
| 2 | 24 | 8 | 31 | 2 | 39 | 4 | 43 |
| 3 | 35 | 9 | 43 | 3 | 33 | 9 | 41 |
| 4 | 27 | 11 | 38 | 4 | 42 | 12 | 54 |
| 5 | 30 | 10 | 39 | 5 | 39 | 6 | 45 |
| 6 | 34 | 9 | 43 |  |  |  |  |
| 7 | 34 | 8 | 42 |  |  |  |  |
| 8 | 33 | 9 | 41 |  |  |  |  |
| 9 | 28 | 11 | 39 |  |  |  |  |
| 10 | 30 | 11 | 41 |  |  |  |  |
| total | 304 | 92 | 396 | total | 184 | 42 | 226 |
| expected<br>distribution if<br>AAbbcc is<br>lethal: 3:1 | 297 | 99 | 396 | expected<br>distribution if<br>AAbbcc is<br>lethal - 4.3:1 | 183.6 | 42.4 | 226 |

### Determination of the CLPC1-TRAP interactome in the absence of CLPF or CLPS1

We extracted the soluble proteome of the three CLPC1-TRAP lines (in WT, *clpf,* or *clps1*) for affinity purification of CLPC1-TRAP to determine its interactome by MSMS and the impact of the loss of CLPS1 or CLPF. Representative plants used for CLPC1-TRAP purifications are shown in Figure 1D. CLPC1-TRAP and its interactome were purified on streptactin columns, and the eluates were run on SDS-PAGE gels, stained with Coomassie BB (Fig. S4). Each gel lane was cut into slices, digested with trypsin, and analyzed by LC-MSMS. We carried out two independent sets of CLPC1-TRAP eluate analyses in WT, *clpf,* and *clps1*, each with three biological replicates (set 1 with replicates 1-3 and set 2 with replicates 4-6). To determine quantitative protein differences between the three genotypes, a minimum observation threshold of 12 AdjSPC across this dataset was applied, identifying 1308 proteins and protein groups, of which 536 were chloroplast-localized proteins (mostly soluble stromal proteins), including 16 out of the 17 chloroplast CLP proteins (not CLPS1) (Dataset S1). There were no significant differences in the total amounts of the CLP protease core (consisting of CLPP, CLPR, and CLPT subunits), indicating that loss of CLPS1 or CLPF did not impact the CLP chaperone-core stability (Fig. 5A). However, CLPF was identified in the WT background in each of the six WT replicates but not in any samples of *clps1* (nor in *clpf,* as expected) (Dataset S1). Figure 5B,C shows the volcano plots for *clps1*/WT and *clpf*/WT. This shows that CLPF is depleted in the CLPC1-TRAP eluates in the *clps1* background, as mentioned above. Immunoblotting detected very low levels of CLPF in the CLPC1-TRAP eluates of the *clps1* background, as compared to the WT background, consistent with the MSMS analysis (Fig. 5D,E). However, no other significant abundance changes were observed for any other chloroplast proteins in the CLPC1-TRAP eluates in *clps1*. In case of the CLPC1-TRAP eluates in *clpf,* the chloroplast ^1^O_2_ sensor EX1 was trapped at a significantly lower level (6-fold) in *clpf* (Fig. 5C). CHLI-1, a subunit of the Mg-chelatase enzyme complex, was also trapped less (3-fold) in *clpf,* although not as much as EX1 (Fig. 5C). Evaluation of the other identified subunits of the Mg-chelatase complex (CHL1-2, CHLD, and CHLH) did not show significant abundant changes (Dataset S1).

**Fig. 5.**
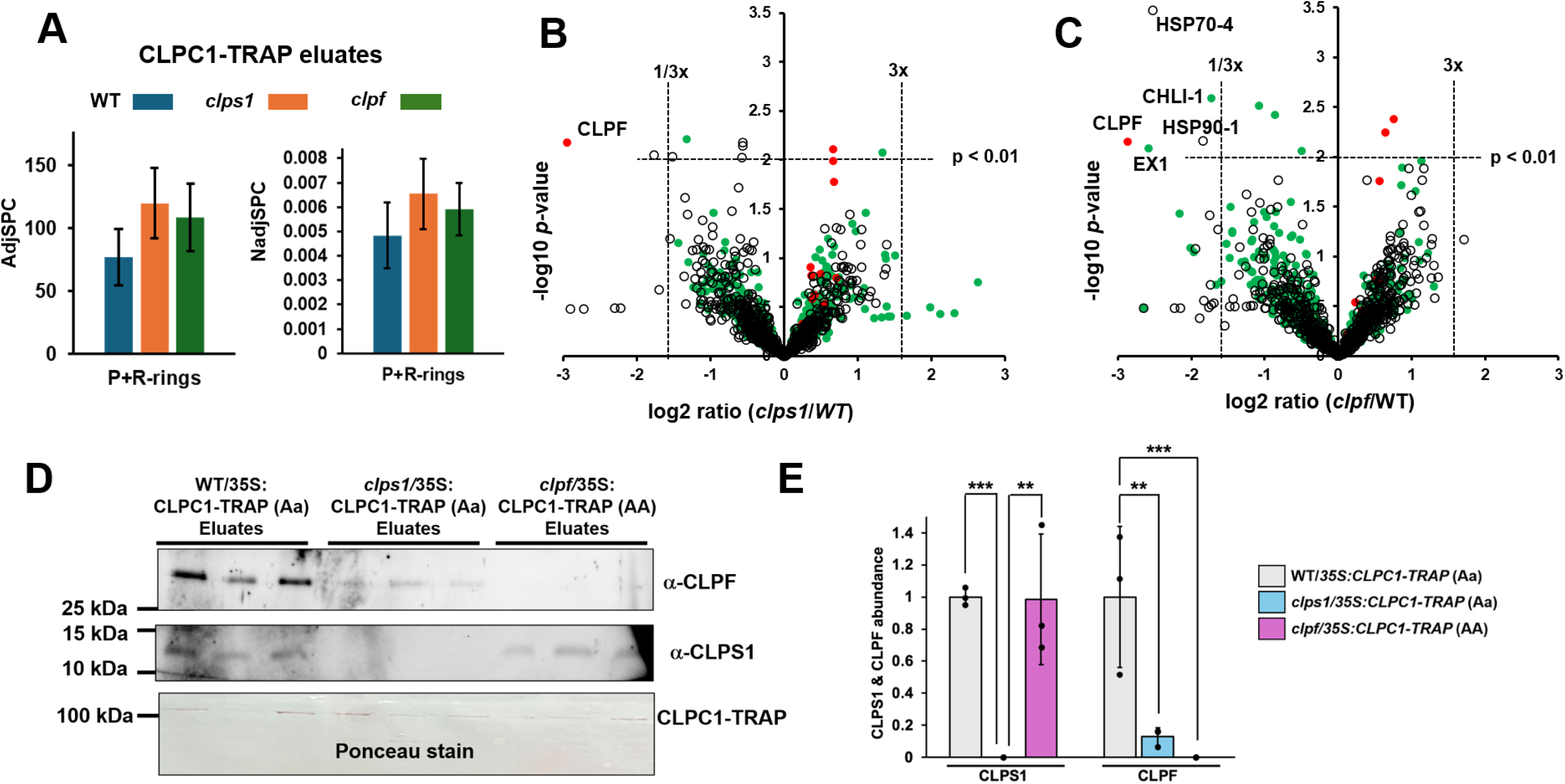
Quantitative MSMS-based proteomics of StrepTactin affinity-enriched CLPC1-TRAP expressed in WT, *clps1,* or *clpf*. **A**. Quantification of identified CLPP and CLPR subunits in the CLPC1-TRAP eluates. Average values (n=6) based on adjSPC and NadjSPC with standard deviations. **B**,**C** Volcano plots of proteins identified and quantified by MSMS in affinity-enriched CLPC1-TRAP of *clpf* and WT (**B**) and *clps1* and WT (**C**). Differentially accumulated proteins (p <0.01) are labelled. Chloroplast proteins are shown in green, and CLP chaperone-protease subunits are shown in red. **D.** Immunoblot of the CLPC1-TRAP eluates of replicates 4-6 for all three genotypes using CLPF- and CLPS1-specific antibodies. A portion of the Ponceau-stained membrane is shown as the loading control. **E.** Quantification of the CLPS1 and CLPF bands from the immunoblot in panel D. Less CLPF is present in the eluates of *clps1*/*35S:CLPC1-TRAP-STREPII* (Aa) than that of WT/*35S:CLPC1-TRAP-STREPII* (Aa), consistent with our MSMS analysis of CLPF in these genotypes. Significance was calculated using Dunnett’s multiple two-way ANOVA comparison. The asterisks indicate significant differences, where ** = p-value < 0.01 and *** = p-value < 0.001. N = 3.

### EX1 protein is stabilized when CLP capacity is reduced in clpr2-1 and CLPC1-TRAP plants

EX1 and its homolog EX2 have been identified as possible CLP substrates, as they are consistently enriched in WT/*35S:CLPC1-TRAP-STREPII* samples when compared to WT/*35S:CLPC1-WT-STREPII* samples. Our current study supports these observations and suggests that the interaction between EX1 and CLPC1-TRAP may be directly or indirectly enhanced by CLPF. To further explore the possibility that EX1 is a CLP substrate, we tested whether the steady-state levels of EX1 and EX2 proteins change in Arabidopsis seedlings of the following eight genotypes: WT, *clpr2-1*, *clpc1-1, clpf*, WT/*35S:CLPC1-WT-STREPII*, and the *35S:CLPC1-TRAP-STREPII* lines in WT, *clpf,* and *clps1* backgrounds (Fig. 6A). Immunoblotting with antibodies specific to EX1 or EX2 showed that the levels of EX1,2 in the *clpf*, *clps1*, and WT/*35S:CLPC1-WT-STREPII* plants were low and comparable to those of WT (Fig. 6B, C). In the *clpc1-1* null mutant, EX1,2 levels were 3-5x higher than in WT (Fig. 6B,C). EX1,2 protein levels were even higher (∼30x) when CLP capacity was very limited by either knockdown of the proteolytic core capabilities in the *clpr2-1* mutant (∼20% of CLP protease capacity remaining (Rudella *et al*., 2006) or by preventing substrate unfolding in the CLPC1-TRAP lines. RT-PCR showed that EX1 mRNA accumulation is the same across these different Arabidopsis lines, demonstrating that EX1 protein abundance is controlled post-translationally (Fig. 6D).

**Fig. 6.**
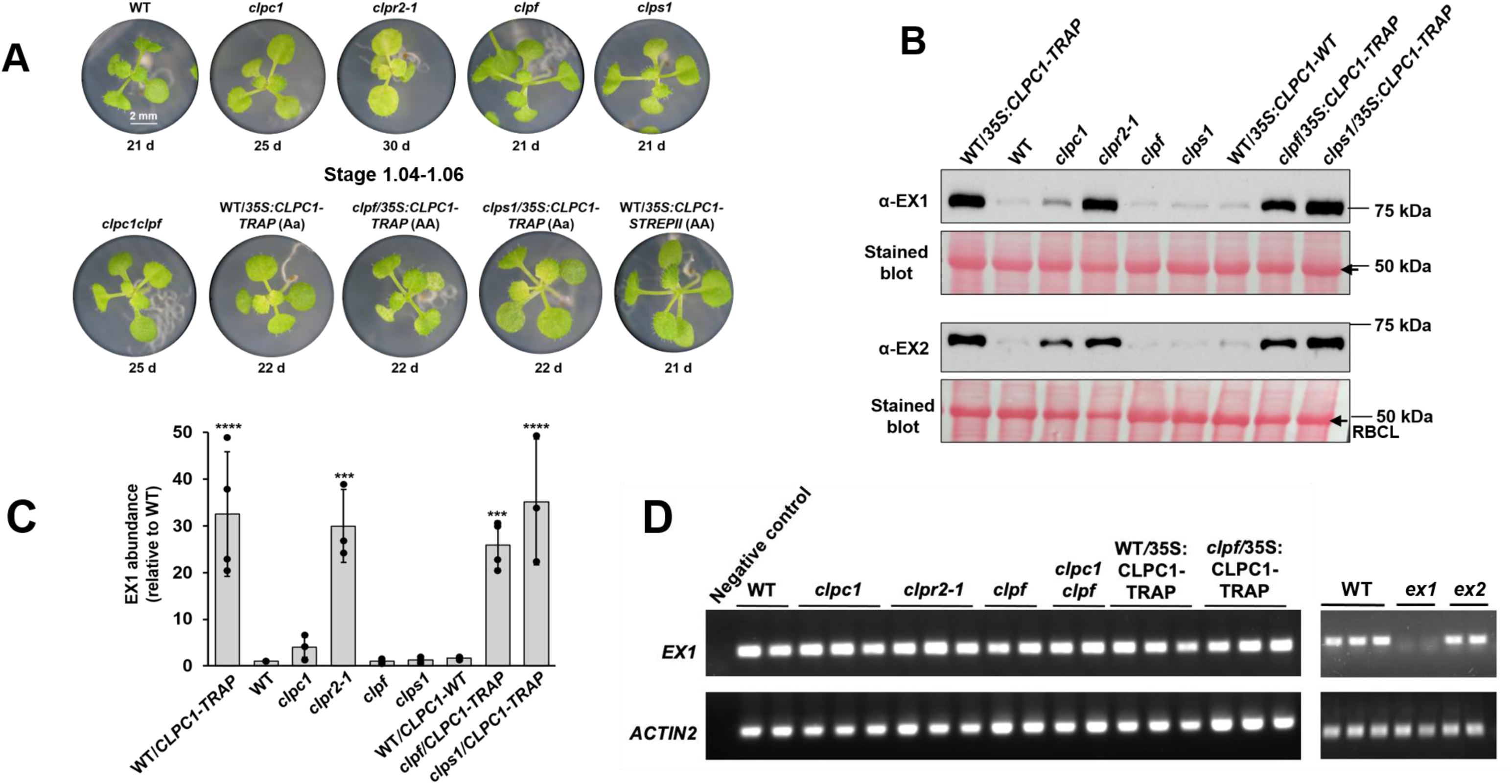
EX1 is stabilized when CLP chaperone or protease capacity is reduced. **A.** Representative images of Arabidopsis plants used for RT-PCR and immunoblot analysis of EX1,2 expression. Seedlings were grown on ½ MS plates (no sucrose) under 10 hr light/14 hr dark conditions until they reached developmental stage 1.04 (2 cotyledons and 4 true leaves). The number of days it took each genotype to reach stage 1.04 (when they were harvested) is indicated. **B.** Immunoblot analysis of EX1 and EX2 accumulation in total (soluble and membrane) leaf proteins extracted from the seedlings shown in panel A. Equal amounts of protein (40 μg) were loaded in each lane and analyzed by immunoblotting with EX1- and EX2-specific antibodies. A portion of the Ponceau-stained membrane is shown as the loading control. The arrow indicates the RUBISCO large subunit **C.** Quantification of the EX1 protein bands from immunoblots. The asterisks indicate significant differences in EX1 levels from those of WT plants, where *** = p-value < 0.001 and **** = p-value < 0.0001. N = 3 or 4. Significance was calculated using Tukey’s one-way ANOVA test. **D.** RT-PCR-based *EX1* mRNA accumulation in the seedlings shown in panel A. *ACTIN2* (*ACT2*) serves as a normalization control for mRNA levels. *EX1* mRNA levels are unchanged in the Arabidopsis plants tested.

### The half-life of EX1 depends on CLPC1 but not CLPF

Our trapping results suggest that the association of EX1 and CLPC1-TRAP relies on the CLPF adaptor. However, EX1 did not accumulate in the *clpf* null background, indicating that EX1 can be degraded in the absence of CLPF. We conducted half-life (CHX-chase) experiments in seedlings of WT and *clpc1* (Fig. 7A) to further test if EX1 is a CLP substrate and whether CLPF modulates the stability of EX1. The half-life of EX1 in WT plants was around seven hours, but significantly longer (>16 hours) in the *clpc1-1* null background (Fig. 7B-D). These data support that the chloroplast CLP system is responsible for EX1 degradation. However, the half-life of EX1 was the same in *clpf* as in WT (Fig. S5), indicating that the degradation of EX1 does not rely on CLPF, in agreement with the steady-state levels of EX1 in *clpf* shown in Figure 6B,C.

**Fig. 7.**
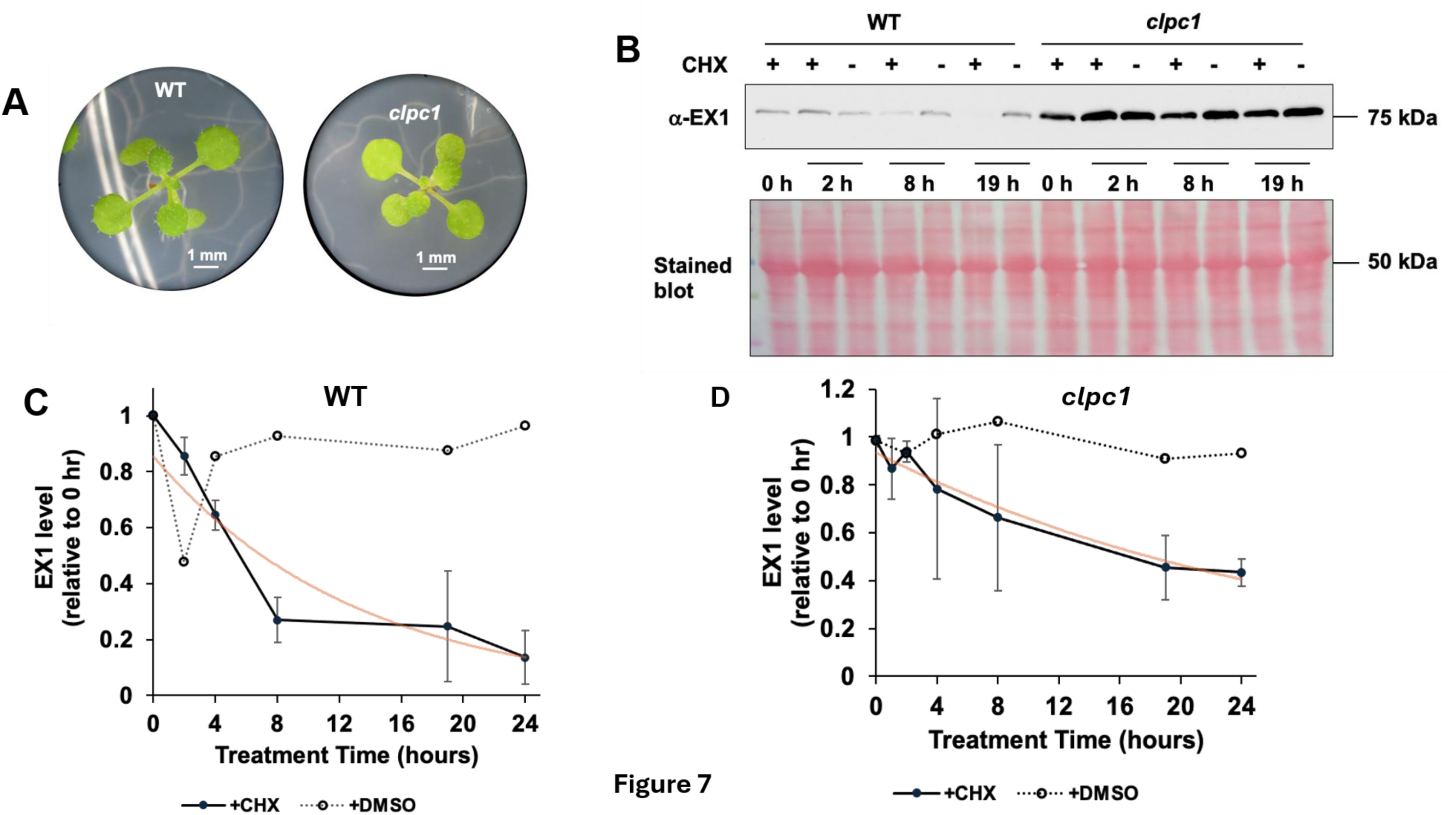
The half-life of EX1 is increased in *clpc1-1* compared to WT. **A.** Representative images of Arabidopsis plants used for the half-life experiments at stage 1.04. Seedlings were grown on ½ MS plates (no sucrose) under 10 hr light/14 hr dark conditions until they reached developmental stage 1.04 (2 cotyledons and 4 true leaves). **B.** Representative immunoblot analysis of EX1 half-life in WT and *clpc1-1* plants. EX1 protein stability was monitored by immunoblotting using an EX1-specific antibody at 0, 2, 8, and 19 hours following treatment with 300 µM CHX in DMSO (+) to inhibit cytoplasmic protein translation. The addition of only DMSO (-) serves as a control. The Ponceau-stained membrane is shown as the loading control (40 µg of total protein). **C, D.** Quantification of the EX1 protein bands in the WT background **(C)** or the *clpc1-1* background **(D)** following either CHX treatment (closed symbols, solid line) or DMSO treatment (open symbols, dashed line). N=3 for the CHX treatment samples, and N=1 for the DMSO control samples. An exponential trend-line for the +CHX data points is shown in orange. For WT, the equation of the line is y = 0.8546e^-0.076x^, R^2^ = 0.907; for clpc1, the equation of the line is y = 0.9342e^-0.035x^, R^2^ = 0.9592. Using these equations, the half-life of EX1 in WT is ∼7 hours, whereas the half-life of EX1 in *clpc1* is ∼18 hours.

## DISCUSSION

### The impact of the loss of CLPS1 and/or CLPF on the CLPC1-TRAP phenotype

The chloroplast CLP chaperone-protease system in Arabidopsis employs substrate-selection mechanisms that involve the CLPS1 and CLPF adaptor proteins (and perhaps additional unknown adaptors) in addition to direct recognition of substrates by the CLP chaperones. CLPS1 is a homolog of the *E. coli* CLPS N-recognin adaptor (Erbse *et al*., 2006; Gao *et al*., 2019) and likely functions as the substrate selector in the chloroplast N-degron pathway (Montandon *et al*., 2019a; Aguilar Lucero *et al*., 2021; Kim *et al*., 2021). The Arabidopsis CLPS1 X-ray crystal structure and *in vitro* affinity assays support that CLPS1 is the functional homolog of bacterial CLPS, albeit with modified N-degron affinities (Colombo *et al*., 2018; Montandon *et al*., 2019a; Aguilar Lucero *et al*., 2021; Kim *et al*., 2021). Arabidopsis CLPS1 interacts with the CLPC1,2 chaperones, CLPF, and several CLP substrates (Nishimura *et al*., 2013; Nishimura *et al*., 2015; Apitz *et al*., 2016). CLPF interacts with CLPS1 independently of the CLPS1 N-degron binding residues (Nishimura *et al*., 2013), and CLPF and CLPS1 act together as a binary adaptor for CLP substrate, GluTR (Nishimura *et al*., 2015). CLPF levels are unchanged in *clps1*, but CLPS1 protein levels increase about 3-fold in *clpf*, indicating either that CLPS1 stability is increased when CLPF is absent or that CLPS1 protein levels are actively upregulated when CLPF is missing (Nishimura *et al*., 2015). Stomal proteomics of *clps1* and *clpf* showed that proteins in both backgrounds generally responded in the same direction, including downregulation of MEP pathway protein 4-hydroxy-3-methylbutyl diphosphate (HDS) and stromal HSP90 and CLPB3, and upregulation of the glycosyl hydrolase DARK INDUCIBLE 1 (DIN1) (Nishimura *et al*., 2015). It is, however, not known whether CLPS1 and CLPF always function cooperatively, such as for GluTR, or whether they can function independently, or even antagonistically (*i.e.,* CLPF or CLPS1 acts as an anti-adaptor) for different CLP substrates. Multi-adaptor systems with two or three adaptors/anti-adaptors have been identified in several bacteria (*e.g.* Caulobacter and Salmonella) and show amazing evolutionary and functional diversity (Mahmoud & Chien, 2018). Hence, an open-ended, discovery-based experimental approach, rather than targeted approaches, could reveal non-canonical functions and pathways for Arabidopsis CLPS1 and CLPF. In this study, we used *in planta* CLPC1-substrate trapping in the presence or absence of CLPS1 or CLPF and follow-up experiments to identify novel *in vivo* substrates of CLPS1 and CLPF.

When expressed in WT Arabidopsis, the *35S:CLPC1-TRAP-STREPII* transgene induces a severe, virescent, photosynthetic dominant-negative phenotype (Montandon et al., 2019). Here we showed that these phenotypes are reproduced in several independent *CLPC1-TRAP-STREPII* lines, thus confirming the causal relationship between accumulation of CLPC1-TRAP in the chloroplast and these phenotypes. The CLPC1-TRAP phenotype is partially suppressed by the loss of *clpf*, but not by the loss of *clps1*, whereas expression of the CLPC1-TRAP transgene results in embryo lethality in the *clps1clpf* double null mutant. These surprising results indicate that the combined loss of CLPS1 and CLPF greatly enhances the negative impact of the CLPC1-TRAP through yet unidentified mechanisms. Loss of CLPS1 and CLPF likely results in increased accumulation of dysfunctional proteins or aggregates either by reduced substrate delivery and/or decreased activity of the remaining endogenous CLPC1 proteins. Surprisingly, whereas loss of CLPS1 does not alter the phenotype of the CLPC1-TRAP line, loss of CLPF suppresses the growth, virescent, and photosynthetic CLPC1-TRAP phenotypes at least in part due to reduced CLPC1-TRAP mRNA and protein accumulation. Collectively, these observations suggest differential contributions of CLPS1 and CLPF to the CLP system. However, loss of both CLPS1 and CLPF has detrimental consequences for viability when the CLP chaperone-protease degradation pathway is severely compromised.

### Loss of CLPF and loss of CLPS1 have differential effects on the pool of CLPC1-trapped proteins

We hypothesized that in the absence of either CLPS1 or CLPF, the pool of CLPC1-trapped substrates would change, reflecting the specific loss of substrate selection by either CLPS1 or CLPF. If CLPF and CLPS1 always function together as a binary adaptor system, then the trapped substrate pools should be the same in the c*lpf* and *clps1* backgrounds. The selected CLPC1-TRAP transgenic lines in the wt, *clpf,* and *clps1* backgrounds accumulated comparable levels of CLPC1-TRAP protein (as determined by immunoblots), allowing comparative CLPC1-TRAP interactome affinity purifications and subsequent MSMS analysis. Most CLPC1-trapped proteins, including the subunits of the CLPPR protease core complex, did not differ significantly across the three genotypes, but there were two statistically significant differences: i) CLPF was trapped at a 10-fold lower level in the *clps1* background, and 2) EX1 was trapped at a six-fold lower level in the *clpf* background. Less trapped CLPF in *clps1* is consistent with our model of binary CLPS1-CLPF adaptor function, resulting in reduced interactions of CLPF with CLPC in the absence of CLPS1. We previously proposed this model based on our *in vitro* binding studies involving recombinant CLPS1, CLPF, GluTR, and several CLPC1 domains (Nishimura *et al*., 2015). The reduced level of trapped EX1 in the *clpf* background might reflect CLPF-enhanced recruitment of EX1 to the CLPC1 chaperone, either with EX1 as a substrate or regulator of CLPC1 through the UVR domains present in CLPF, EX1, and CLPC1, as we recently discussed (Annis *et al*., 2024).

### The CLP system controls steady-state levels of EX1,2

The results of our trapping experiments led us to hypothesize that EX1 is a CLP substrate and that EX1 might rely on the CLPF adaptor for its degradation. In our previous CLPC1 trapping studies, we showed that EX1,2 are very abundant in the CLPC1 trap eluates in the WT background, especially given that EX1,2 proteins accumulate at only very low levels in WT plants, as illustrated by results in the Arabidopsis PeptideAtlas (Montandon *et al*., 2019b; Rei Liao *et al*., 2022). In this study, we showed by immunoblotting of total leaf protein extracts from a range of genotypes that both EX1 and EX2 protein levels increased more than 30-fold in the *clpr2-1* mutant and also in the CLPC1-TRAP lines in WT, *clpf,* and *clps1* backgrounds. EX1,2 levels were also several-fold higher in the *clpc1* null line; this increase was not as dramatic as for the CLPC1-TRAP lines and *clpr2-1* line, likely because the closely related homolog CLPC2 is upregulated in *clpc1* and partially complements substrate unfolding and delivery capacity (Constan *et al*., 2004; Sjogren *et al*., 2004; Kovacheva *et al*., 2007; Nishimura *et al*., 2013). This dramatic increase of EX1 and EX2 was not due to transcriptional upregulation but instead due to protein stabilization. Our *in vivo* protein experiments demonstrated that the half-life of EX1 is 3 times longer in *clpc1* compared to WT (∼7 hrs vs ∼18 hrs). Interestingly, whereas EX1 was trapped significantly less in the *clpf* background, it was not stabilized in *clpf.* Consistent with the EX1 level in *clpf,* the half-life of EX1 was similar in WT and *clpf,* indicating that EX1 degradation by CLP does not strictly require CLPF. The apparent discrepancy between the CLPC1 trapping results in *clpf* and the lack of impact on half-life and steady-state level of EX1 in *clpf* remains to be determined.

Collectively, our data and previous publications (Montandon *et al*., 2019b; Rei Liao *et al*., 2022) strongly support a model in which the CLP chaperone-protease system controls the steady-state levels of EX,1,2 upstream of the EX1,2 ^1^O_2_ sensing and oxidation and subsequent FTSH-driven degradation and retrograde signaling (Fig. 8). Pull-down assays with EX1-GFP by the Kim lab identified CLPC1 among interacting proteins (Dogra *et al*., 2019), which is consistent with our results. Figure 8 depicts our straightforward working model of EX1,2 homeostasis: As nuclear-encoded proteins, EX1 and EX2 are transcribed in the nucleus, then translated into pre-proteins containing N-terminal chloroplast transit peptides in the cytosol. These pre-proteins are imported into the chloroplast, where they are processed by stromal processing peptidase. Mature EX1 and EX2 are then sorted to the stroma-facing side of the thylakoid grana margins, where they sense and respond to ^1^O_2,_ resulting in their specific oxidation and thylakoid FTSH-mediated degradation. Our results suggest that the majority of mature EX1,2 does not accumulate at the thylakoid membrane but is instead degraded by the CLP chaperone-protease system.

**Fig. 8.**
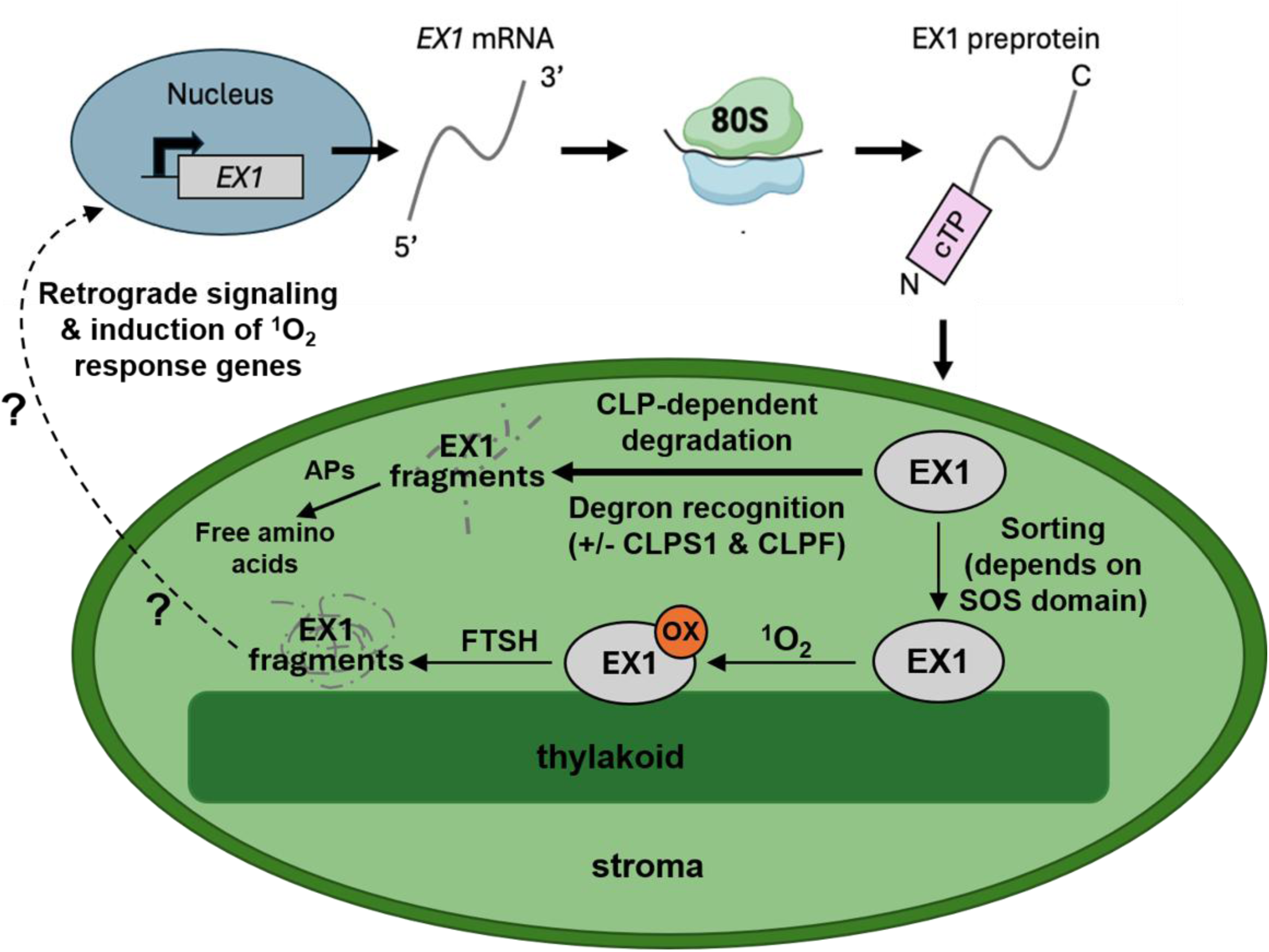
Working model of EX1 protein homeostasis. Following translation in the cytoplasm, EX1,2 preproteins are imported into the chloroplast. The CLP chaperone-protease system continuously degrades a large percentage of imported EX1,2 protein to maintain a low, basal level in the chloroplast. APs are amino peptidases that can degrade small peptides that are released by the CLP protease. EX1,2 associates with the thylakoid membrane via its SOS domain. At the thylakoid membrane, EX1 interacts with singlet oxygen (^1^O_2_), which causes oxidation of W643 located in the SOS domain. This tryptophan oxidation triggers EX1,2 degradation by FTSH followed by downstream retrograde signaling through unknown molecular players. See (Lee & Kim, 2024) for details and references on EX1,2 functions. There are conflicting reports of extra-chloroplast location and function of EX1 (Liu *et al*., 2024) (Li *et al*., 2023; Zhao *et al*., 2025), which are not considered in our working model, given their uncertain nature.

Future experiments should provide a better understanding of why EX1,2 protein levels in the chloroplast are maintained at such low levels. However, there have been several other observations where the steady-state levels of low-abundant chloroplast proteins such as GUN1 and DUF760-1 are also controlled by the CLP protease system (Wu *et al*., 2018; Yuan & van Wijk, 2024). These proteins are transcribed and translated, but then quickly turned over through proteolysis by CLP. Comparison of large-scale public Arabidopsis mRNA data (https://bar.utoronto.ca/) and mass spectrometry-based protein data (https://peptideatlas.org/builds/arabidopsis/) identified additional chloroplast proteins with low abundance protein levels (either not detected by MSMS or only with very few observations) but high mRNA levels such as several of the chloroplast sigma factors (*e.g*. SIG1,3), suggesting additional chloroplast proteins that are maintained at low levels through high protein turnover rates (van Wijk *et al*., 2024). Collectively, this shows that chloroplast proteostasis is poorly understood and warrants future investigation.

## Acknowledgments

This research was supported by the National Science Foundation grant MCB-2322813 to K.J.v.W. C.M.R. was also supported by a Chemistry-Biology Interface Training grant (National Institute of Health/National Institute of General Medical Sciences grant T32GM138826). We thank Marissa Annis for discussions and support.

## Data availability

The data generated in this study are included in this article and the online supplementary material.

## Author Contributions

C.M.R, P.R., B.Y., I.B., M.M.J and K.J.V.W. conceived, designed, and carried out experiments. All authors analyzed the results. C.M.R and K.J.V.W. wrote the article. All authors read and contributed to the final article.

## Supporting Information

**Fig. S1.**
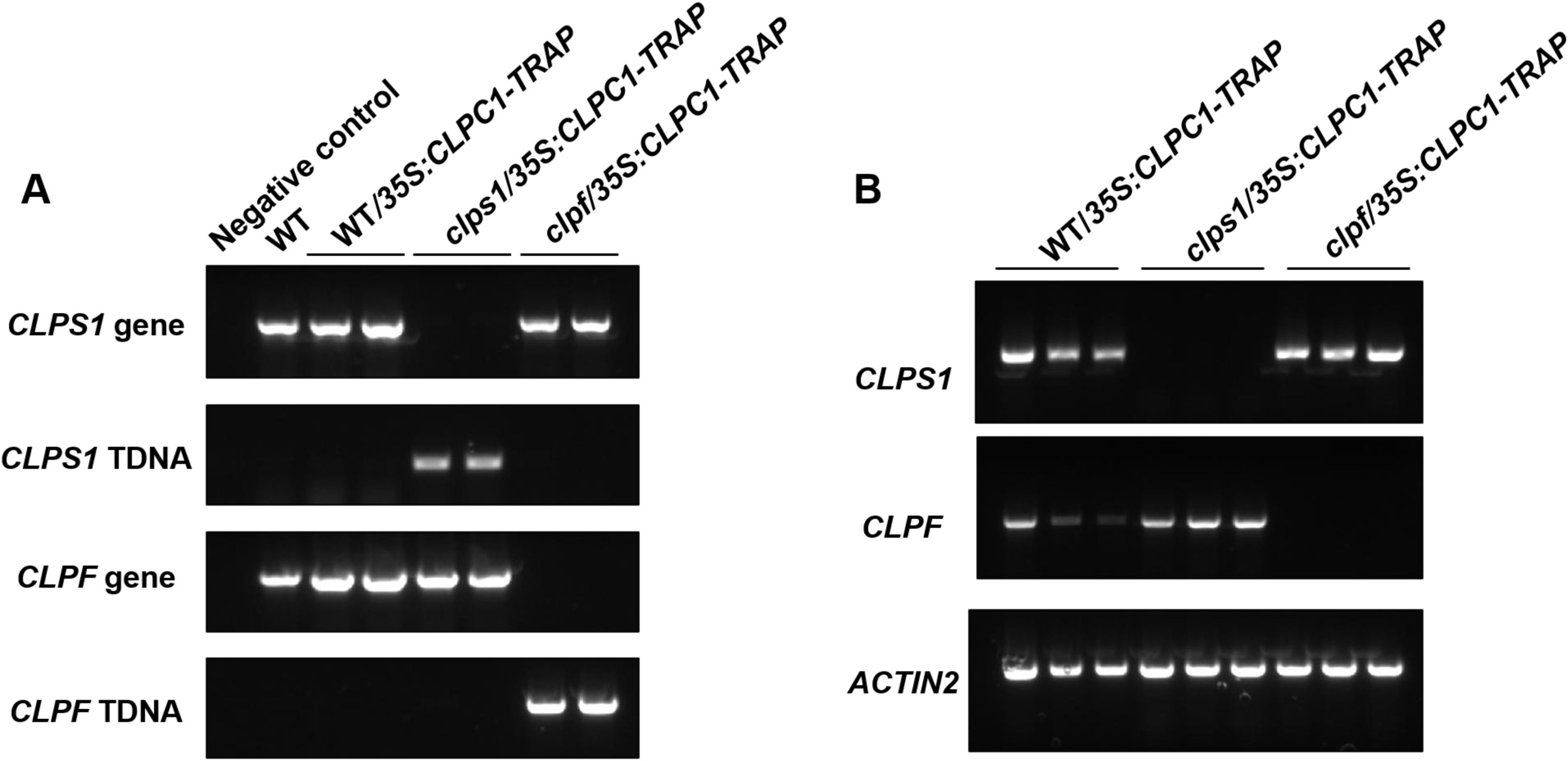
Genotyping of the clpf/35S:CLPC1-TRAP-STREPII, clps1/35S:CLPC1-TRAP-STREPII, and WT/35S:CLPC1-TRAP-STREPII lines. **A.** PCR (30 cycles) of genomic DNA extracted from WT, WT/35S:CLPC1-TRAP-STREPII (Aa), clps1/35S:CLPC1-TRAP-STREPII (Aa), and clpf/35S:CLPC1-TRAP-STREPII. (AA) plants. **B.** RT-PCR (25 cycles) of cDNA made from RNA extracted from WT/35S:CLPC1-TRAP-STREPII (Aa), clps1/35S:CLPC1-TRAP-STREPII (Aa), and clpf/35S:CLPC1-TRAP. (AA) plants. ACTIN2 is used as a loading control.

**Fig. S2.**
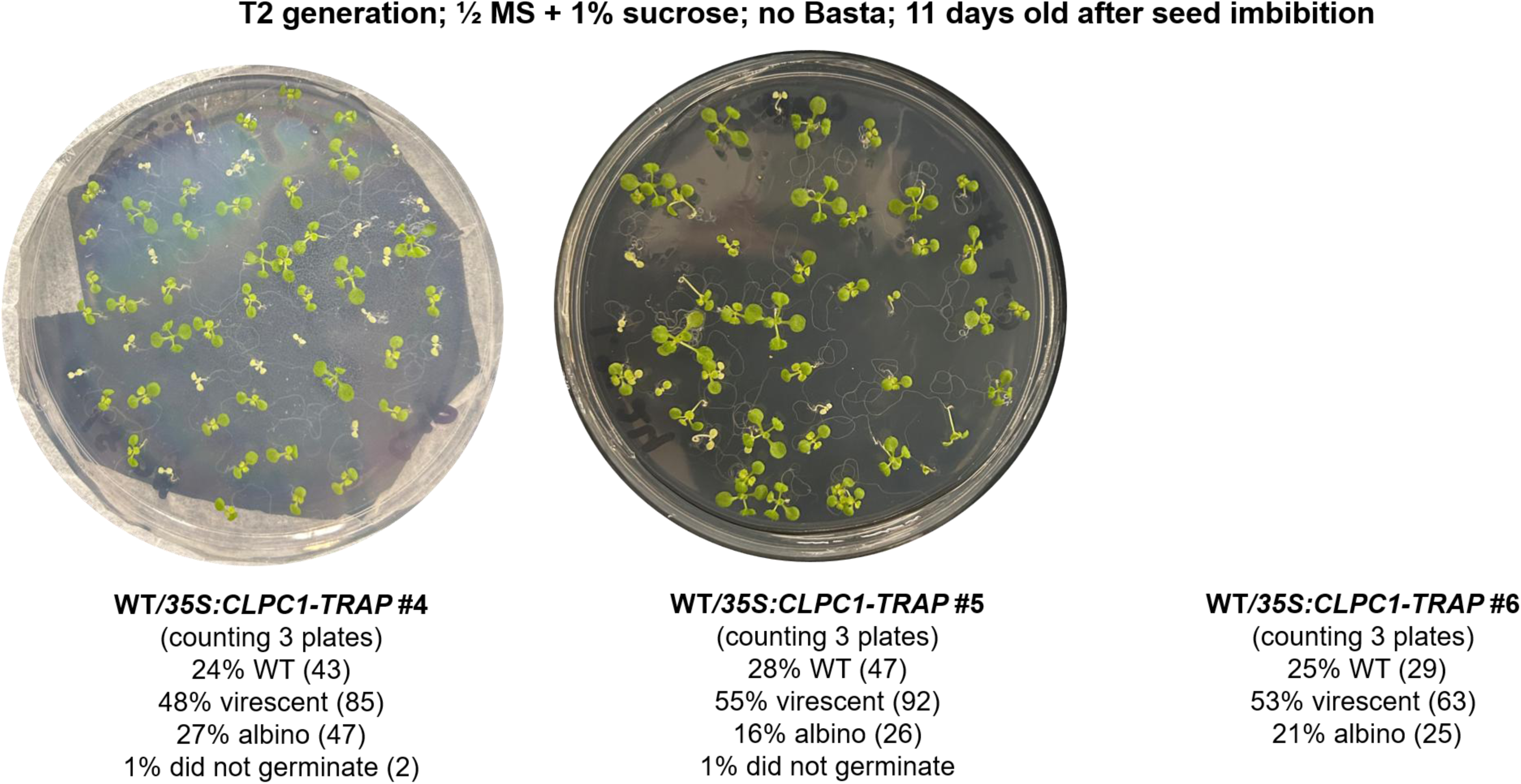
Generation of three independent WT/*35S:CLPC1-TRAP-STREPII* lines. T2 generation of three additional, independent WT/*35S:CLPC1-TRAP-STREPII* lines grown without selection media (no Basta). A total of 3 plates were grown for each of the three lines (#4,5,6), and seedlings were counted for phenotypes, distinguishing between WT, virescent, albino, and not germinated.

**Fig. S3.**
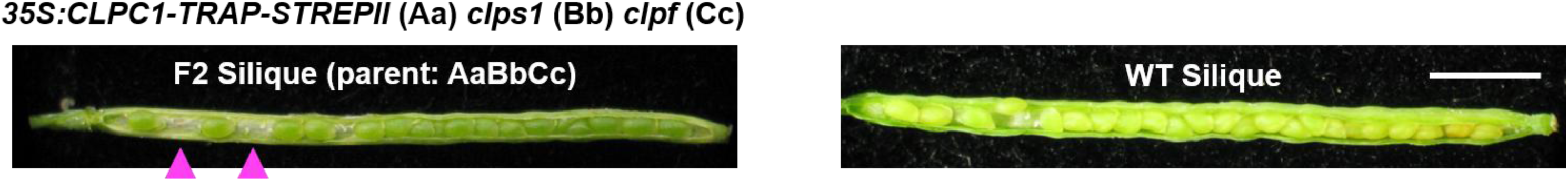
Representative images of developing siliques from a WT plant and an F2 plant (from the cross in Fig. 1A) heterozygous for *clps1* (Bb), *clpf* (Cc), and *35S:CLPC1-TRAP-STREPII* (Aa). The F2 silique shows embryo abortions indicated by arrows. Scale bar = 1 mm.

**Fig. S4.**
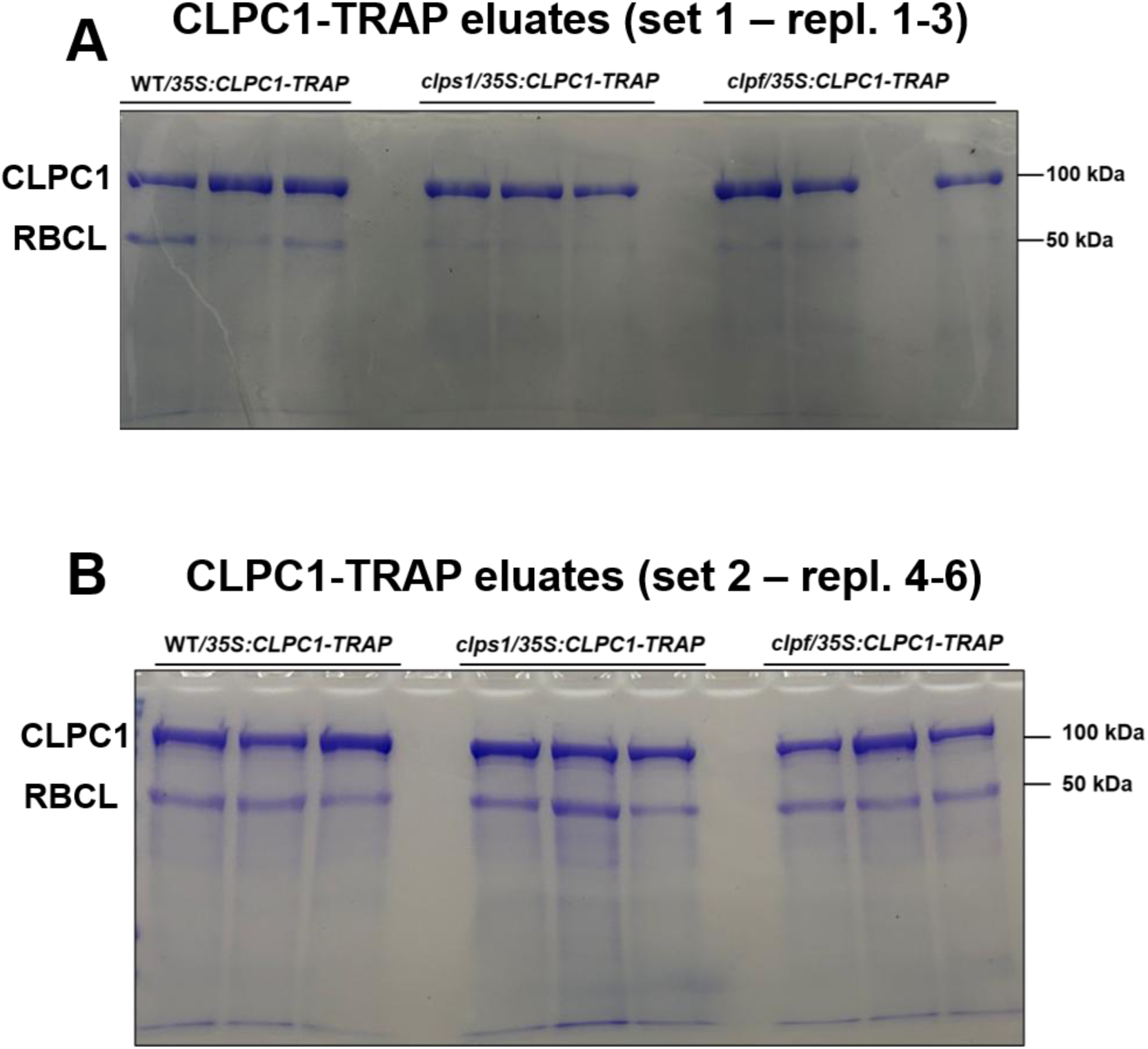
SDS-PAGE gels of streptactin eluates for MSMS analysis. **A,B.** SDS-PAGE of StrepTactin-purified samples from set 1 (replicates 1-3) (**A**) and set 2 (replicates 4-6) (**B**).

**Fig. S5.**
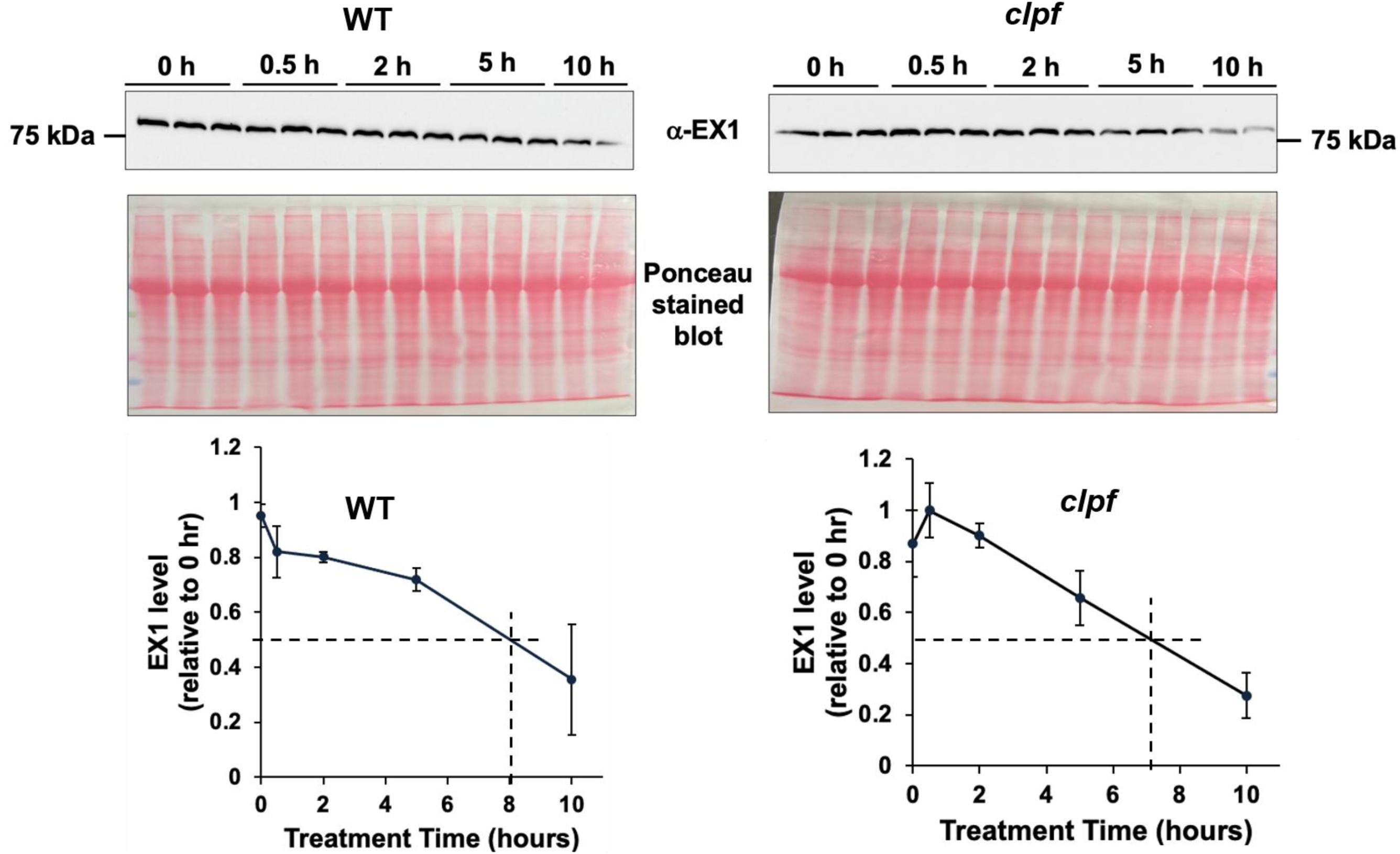
Half-life determination of EX1 in WT and *clpf* seedlings. EX1 protein accumulation and turnover were monitored by immunoblotting using an EX1-specific antibody at 0, 0.5, 2, 5, and 10 hours following treatment with 300 µM cycloheximide (CHX) to inhibit cytoplasmic protein translation. The Ponceau-stained membrane is shown as the loading control (40 µg of total protein). The half-life of EX1 in both the WT and *clpf* background is ∼7-8 hours.

**Table S1.**
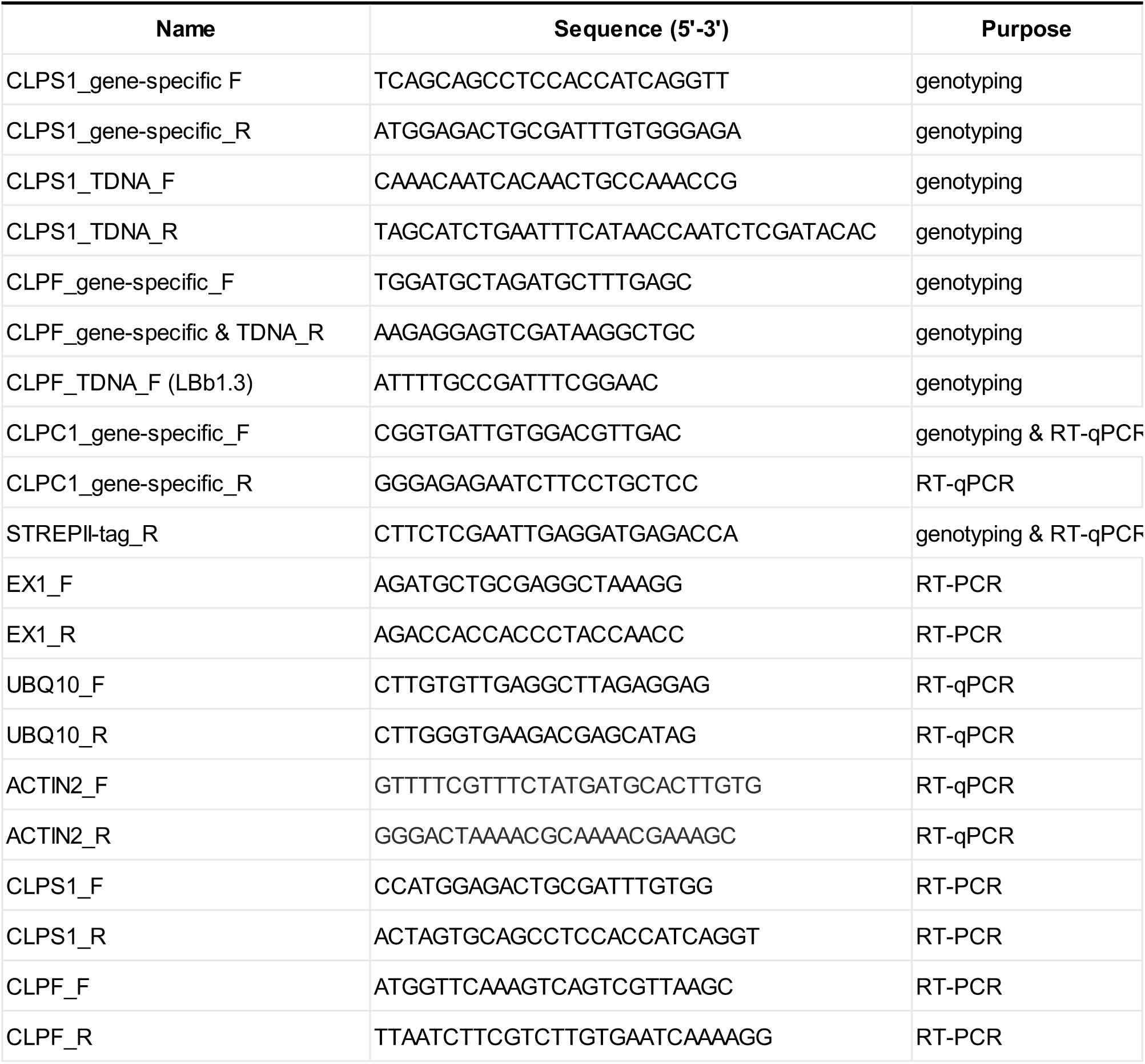
Primers used for genotyping and RT-(q)PCR in this study.

**Dataset S1.** MSMS-based proteome analysis of streptactin eluates of WT/*35S:CLPC1-TRAP-STREPII*, *clpf/35S:CLPC1-TRAP-STREPII* and *clps1/35S-CLPC1-TRAP-STREPII*.

